# RELB Reprograms Exhausted Tumor-Infiltrating Lymphocytes for Improved Adoptive Cell Therapy

**DOI:** 10.1101/2025.10.11.681829

**Authors:** Christian D. McRoberts Amador, Rachel E. Conover, Michael C. Brown, Liliana S. Lyniv, Pamela K. Noldner, Ying Zhou, Aretha R. Gao, Sean R. McCutcheon, Scott J. Antonia, Charles A. Gersbach

## Abstract

Tumor-infiltrating lymphocytes (TILs) are a promising autologous cell therapy to treat solid tumors. TILs are manufactured by expanding and reinfusing tumor-reactive T cells from tumor biopsies. Efficacy of TIL therapies has been limited by the heterogeneity of expanded TIL products and the high prevalence of dysfunctional exhausted CD8+ T cells (T_EX_). While a subset of CD8+ TILs co-expressing CD103 and CD39 are enriched for tumor-reactive TILs across multiple cancer types, these cells are often in the T_EX_ state with low proliferative potential. To identify regulators of human TIL proliferation, we screened an open reading frame library encoding for all human transcription factors (TFs). RELB emerged as the dominant driver of human TIL expansion with a skew towards CD8+ cells. TCR diversity was maintained after multiple days of *in vitro* expansion driven by RELB. Transcriptome profiling of multiple RELB-expressing TIL subtypes revealed a shift towards a memory/costimulatory-like phenotype. Using a HER2-targeting CAR and tumor co-culture model, RELB conferred improved persistence after multiple tumor challenges *in vitro* and improved solid tumor control in mouse xenografts *in vivo*. Finally, co-culture of RELB-overexpressing TILs with patient-matched tumor organoids showed an increase in TIL product polyfunctionality, tumor reactivity, and tumor killing. Collectively these results support promoting RELB expression as a strategy for broadly enabling TIL therapy for treating solid tumors.

## Introduction

Adoptive T cell therapy (ACT) has revolutionized cancer treatment by employing effector T cells to directly kill cancer cells. Chimeric antigen receptor (CAR)-engineered T cell therapies have had incredible success in blood cancer indications, leading to durable responses and FDA approval of seven autologous CAR-T products^1-7^. Tumor-infiltrating lymphocytes (TILs) are a promising autologous T cell therapy to treat solid tumors by expanding and reinfusing polyclonal, tumor-reactive T cells isolated from primary tumor resections in an antigen agnostic manner^8,9^. While TIL therapy has had some clinical success in multiple cancer indications^10-15^, including recent FDA approval for advanced metastatic melanoma^16-18^, its efficacy has been limited by the heterogeneity of expanded TIL products and the high prevalence of dysfunctional terminally-exhausted T cells (T_EX_)^19-23^. The activity of these T_EX_ cells is essential for response to TIL therapy since they are the TIL subset reactive to tumor-specific antigens^23,24^, but they are predominantly characterized by impaired proliferative and cytotoxic capabilities^25^.

A subset of double-positive (DP) CD8+ TILs co-expressing CD103 and CD39 surface proteins are enriched for tumor-reactive tissue-resident TILs across multiple cancer types^26-28^. These DP TILs effectively kill matched tumors, but still display hallmarks of T cell exhaustion and eventual disfunction^28^. While these DP TILs are enriched for tumor-reactive TILs that are functional *in vitro*, a more stem-like CD8+ TIL phenotype has been associated with durable clinical response^29-31^.

High-throughput genetic screens in primary human T cells from peripheral blood have elucidated numerous aspects of T cell exhaustion biology, with several gain/loss-of-function screens identifying factors that improve T cell phenotype for CAR-T cell therapy^32-36^. Several screening efforts have focused on the prevention of T cell exhaustion^32,37^, but factors that mediate reversal of an exhausted phenotype in primary human T cells have not been identified. Screening in primary human TILs presents a unique opportunity to discover factors that reverse T cell exhaustion.

To systematically identify factors that reinvigorate T_EX_ cells via cell state reprogramming, we screened a lentiviral Open Reading Frame (ORF) library encoding for all human transcription factors (TFs)^38^ in tumor-reactive human DP TILs from non-small cell lung cancer patients (NSCLC). TFs associated with cell proliferation (MYC, MYC-L, and RELA) emerged as top hits for the CD8+ TIL screen, but only RELB emerged as a hit driving expansion of DP TILs. RELB overexpression led to improvement in TIL proliferation, transcriptional reprogramming of DP TILs towards a memory and costimulatory-like state, and enhanced *in vitro* and *in vivo* tumor control.

## Results

### Proliferation screen in exhausted CD103+ CD39+ human NSCLC TILs identifies RELB as a positive regulator of CD8+ TIL expansion

To screen for factors that improve proliferation of exhausted TILs, we used a library of full-length ORFs of all human TFs and related isoforms^38^. Bulk CD8+ or enriched CD8+ DP TILs were transduced with the library of TF ORFs, labelled with CellTrace to assess cell proliferation, and stimulated with anti-CD3/CD28 in vitro (Fig. 1A, Supplemental Fig. 1A-D). Highly proliferative (CellTrace low) and lowly proliferative (CellTrace high) TILs were isolated by fluorescence activated cell sorting (FACS) and barcode distributions were sequenced to identify candidate TFs that selectively improve DP TIL expansion relative to other CD8+ subtypes (Fig. 1A, Supplemental Fig. 2A-B).

**Fig. 1:**
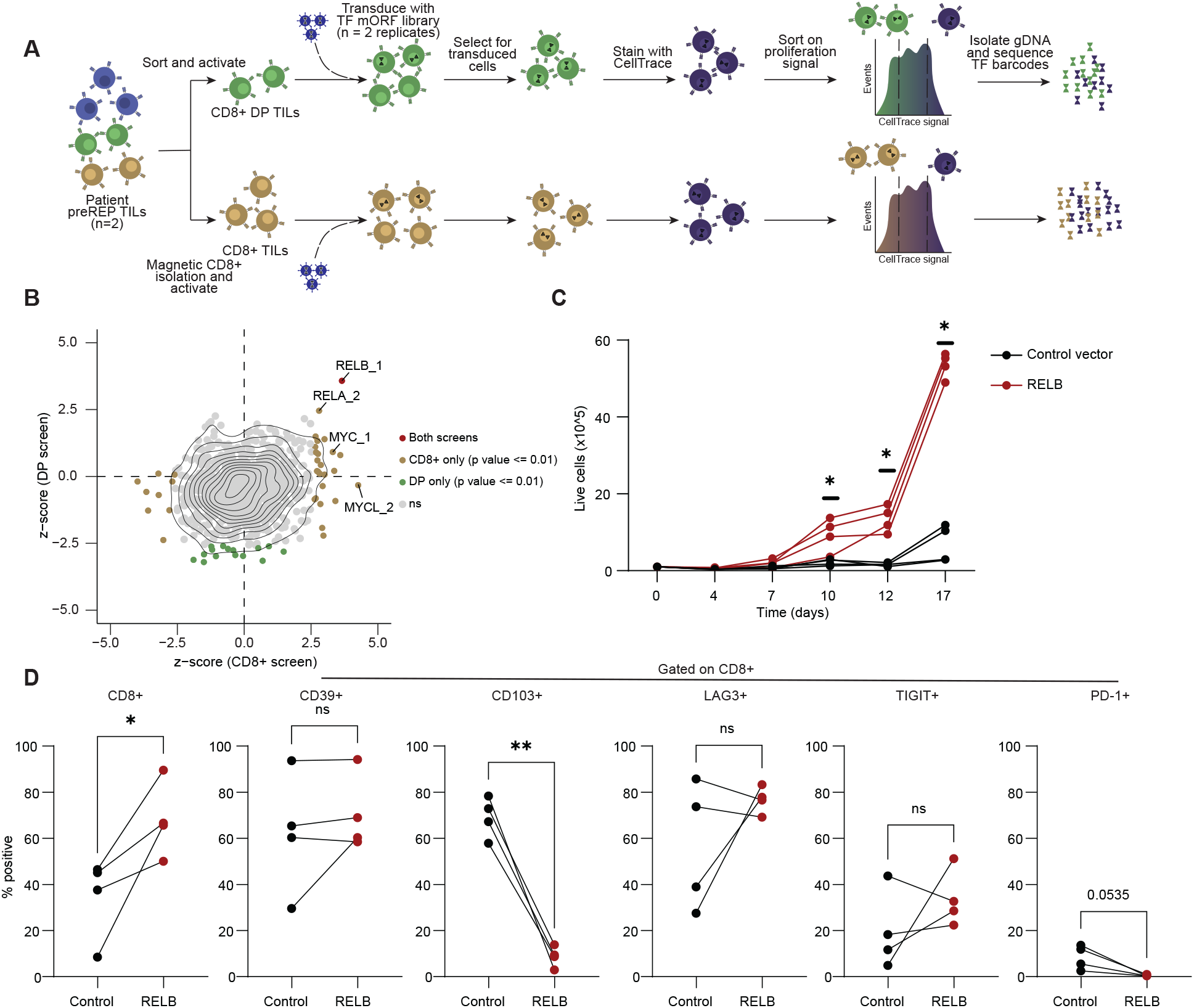
Proliferation-based FACS screen in CD8+ and CD8+ DP TILs reveals TF regulators of proliferation of primary human NSCLC TILs. A) Experimental layout describing FACS-based human NSCLC TIL screen for regulators of proliferation in CD8+ and CD8+ DP TILs. B) Z-scores of DP screen versus z-scores of CD8+ TILs from DESeq2 analysis. To ensure robust coverage of each TF, TF barcodes with less than 500 mean counts were filtered out. ORF enrichment was defined using a paired two-tailed DESeq2 test with Benjamini–Hochberg correction and using a p. value cutoff of < 0.01. C) Proliferation validation in four distinct NSCLC preREP TIL patient samples for RELB ORF overexpression versus Thy1.1 ORF control overexpression. D) Flow cytometry analysis at the end of proliferation experiment for T cell lineage (CD8), tissue residency (CD103), and exhaustion (CD39, LAG3, PD-1, TIGIT) markers.

Several canonical positive regulators of cell proliferation (*MYC, MYC-L, RELA*) improved CD8+ T cell proliferation in this screen, but only *RELB* emerged as a strong hit for DP TIL proliferation (Fig. 1B, Supplemental Table 1, Supplemental Data). RELB is a key cofactor responsible for non-canonical NF-κB signaling^39^. RELA-mediated canonical NF-κB signaling is driven by T cell receptor (TCR) stimulation to drive T cell proliferation^40^, while RELB-mediated non-canonical NF-κB signaling is driven by co-stimulation of TCRs with other T cell surface receptors and is not required for T cell activation^40,41^. Interestingly, non-canonical NF-κB signaling is associated with long-lasting memory T cell formation and can be activated via signaling of costimulatory molecules belonging to the TNFRSF family that are upregulated after initial TCR activation^39,40,42^.

Transduction of unsorted TILs with RELB lentivirus revealed a significant increase in proliferation when compared to control vector (Fig. 1C). We also observed enhanced proliferation in PBMC-derived CD8+ T cells (Supplemental Fig. 3A), indicating the effect of RELB may not be specific to the exhausted state. Flow cytometry analysis at day 17 post-transduction revealed a significant increase in proportion of CD8+ TILs (Fig. 1D). We also tested a different RELB-expressing lentiviral construct that co-expresses GFP as a reporter rather than a puromycin selection cassette and confirmed an increase in CD8+ proportions in all four TIL patient samples tested (Supplemental Fig. 3B). Intriguingly, there was a decrease in expression of CD103 and PD1 in the expanded RELB-expressing CD8+ TILs (Fig. 1D), suggesting a transition away from tissue residency towards a less exhausted state.

Importantly, this proliferative state was dependent on TCR stimulation since TILs that were transduced with RELB+GFP reporter lentivirus without anti-CD3/CD28 stimulation did not show TCR-independent growth (Supplemental Fig. 3C-D). Cytokine-independent proliferation was also not observed, as IL-2 starvation eventually halted RELB-mediated growth (Supplemental Fig. 3E).

Since sustained overexpression of RELB from an integrated lentiviral vector may not be desirable for clinical use, we investigated if transient RELB expression could similarly shift the phenotype of exhausted TILs. We achieved high mRNA electroporation efficiency in primary human TILs of ∼50-80% (Supplemental Fig. 4A). Transient delivery of RELB via mRNA electroporation early in expansion (2 days post-activation) also demonstrated a skew towards CD8+ TILs at seven days post-electroporation (Supplemental Fig. 4B), and electroporation of RELB mRNA later in expansion (day 16 post initial activation) conferred improved proliferation in three out of four patient samples tested (Supplemental Fig. 4C). This improvement in proliferation was seen in both CD4+ and CD8+ TILs (Supplemental Fig. 4D-F).

### RELB maintains TCR diversity in expanded TIL products and reprograms the transcriptome of exhausted CD8+ CD103+ CD39+ TILs towards a memory/costimulatory-like state

We next tested the impact of RELB overexpression on T cell receptor (TCR) diversity and transcriptome by sorting either CD4+ or CD8+ non-DP TILs (i.e. CD103+ CD39-, CD103-CD39+, and CD103-CD39-), and CD8+ DP TILs transduced with either Thy1.1 control vector or RELB lentivirus, both of which co-express the GFP reporter (Fig. 2A, Supplemental Fig. 5A). RELB overexpression did not impact TCR diversity as measured by the effective number of TCR clones represented in the expanded cell population (Fig. 2B). Strikingly, RELB overexpression led to robust expansion of CD8+ DP TIL for three out of four patient samples, whereas none of the samples treated with control vector expanded sufficiently for analysis; only two patient samples expanded sufficiently for the untransduced control conditions (Fig. 2B, right panel). RELB also conferred a significant improvement in proliferation for both CD4+ and CD8+ non-DP TIL subsets across all patient samples tested (Supplemental Fig. 5B-C), and CD8+ TILs yielded the highest increase in proliferation (Supplemental Fig. 5B). DP TILs had the highest mean percent of GFP+ cells for all RELB conditions at the end of expansion (Supplemental Fig. 5C).

**Fig. 2:**
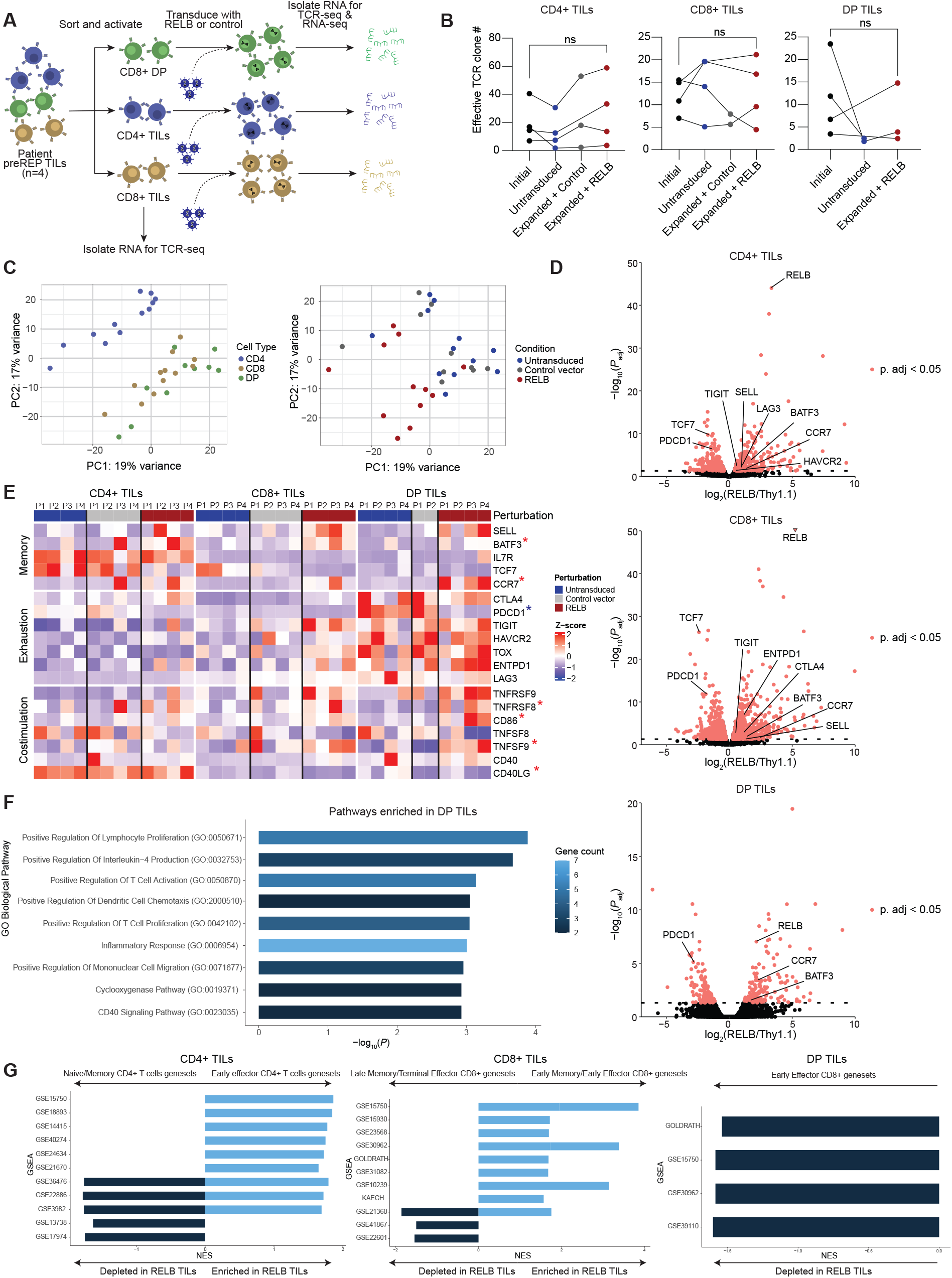
RELB maintains TCR repertoire diversity, pushes DP TILs to a more memory/costimulatory-like state, and partially reverses exhaustion. A) Experimental layout describing transcriptomic and T cell receptor diversity profiling via bulk RNA-seq and TCR-seq on CD4+, CD8+, and CD8+ DP NSCLC TIL subsets. B) Effective TCR clone number in initial (day 0) and expanded (day 19). Expanded conditions were either not transduced with lentivirus or transduced with corresponding GFP+ORF (Thy1.1 or RELB) lentivirus 24 hours post-activation. P values were calculated using paired t-tests. (C) Principal component analysis (PCA) plots for all RNA-seq samples analyzed with DESeq2. (D) Significance (p_adj_) versus fold change between RELB and Thy1.1 samples in different TIL subtypes. Gene enrichment was defined using a paired two-tailed DESeq2 test with Benjamini–Hochberg correction and a p_adj_ cutoff of <0.05. (E) Heatmap of relevant genes related to T cell memory, exhaustion, and costimulation. Each column represents a different patient sample within a specified perturbation. Genes asterisked in red are differentially enriched in DP TILs overexpressing RELB when compared to Thy1.1, while genes asterisked in blue are differentially depleted in DP TILs overexpressing RELB when compared to Thy1.1. (F) GO Biological pathway enrichment analysis on differentially enriched pathways (p < 0.05) in DP TILs. (G) Relevant GSEA enriched and depleted gene sets in CD4+, CD8+, and DP TILs for RELB vs Thy1.1 comparison (p_adj_ < 0.05). Positive Normalized Enrichment Score (NES) corresponds to enrichment of pathway in RELB-overexpressing conditions while a negative NES corresponds to depletion.

We also assessed the transcriptomic changes RELB was driving in each subset by performing bulk RNA-seq of treated cells. As expected, principal component analysis of the RNA-seq datasets revealed co-clustering of CD4 TIL samples and CD8 with DP TIL samples (Fig. 2C). RELB overexpressing samples co-clustered together in the CD8 and DP TIL samples (Fig. 2C). Differential analysis of RELB overexpression versus control vector revealed hundreds of differentially expressed genes (DEGs) in all TIL subsets (Fig. 2D, Supplemental Table 2). Of note, only two Thy1.1 control vector DP populations sufficiently expanded for RNA isolation, underpowering the DP RELB vs Thy1.1 comparison relative to the CD4+ and CD8+ non-DP comparisons. The transcriptomic changes driven by RELB positively correlated across all TIL subpopulations (Supplemental Fig. 6A). As expected, untransduced and control vector DP TIL conditions were defined by an exhausted state with high levels of expression of *CTLA4, PDCD1, TIGIT, HAVCR2, TOX, ENTPD1*, and *LAG3* when compared to CD4+ and CD8+ TILs (Fig. 2E). Strikingly, RELB reprogrammed exhausted DP TILs towards a more memory and costimulatory-like state with upregulation of key T cell memory genes (*SELL, BATF3, CCR7*), significant downregulation of exhaustion marker *PDCD1*, and upregulation of several costimulatory receptors and ligands (*TNFRSF9, TNFRSF8, CD86, TNFSF9, CD40LG*) (Fig. 2E, Supplemental Table 2).

Biological pathway enrichment analysis of genes that were differentially enriched with RELB overexpression in DP TILs versus vector control revealed an enrichment of lymphocyte proliferation, T cell activation, IL-4 production, and CD40 signaling pathways (Fig. 2F, Supplemental Table 3). Gene-set enrichment analysis revealed multiple significantly enriched datasets in all TIL subsets (Fig. 2G, Supplemental Fig. 6B-D, Supplemental Table 3). Overall, CD4+ cells overexpressing RELB enriched in early CD4+ effector gene sets while CD4+ naïve/memory gene sets were depleted, CD8+ cells overexpressing RELB enriched in CD8+ early memory/effector gene sets while CD8+ late memory/terminal effector gene sets were depleted, and DP cells overexpressing RELB depleted CD8+ early effector gene sets (Fig. 2G, Supplemental Table 3). Collectively, the results suggest that RELB overexpression pushes CD4 and CD8 TIL subtypes towards an early effector-like state, while DP TILs are pushed towards memory/costimulatory-like state and away from an effector state.

### RELB confers improved persistence to CD8+ TILs in a rechallenge tumor killing assay

To assess whether TIL cytotoxic capabilities and persistence were impacted by RELB, we conducted an *in vitro* tumor rechallenge assay by arming TILs with a HER2-targeting CAR, overexpressing RELB, and co-culturing with HER2+ SKBR3 breast cancer cells expressing nuclear GFP (Fig. 3A). The first two challenges did not show a difference in tumor control with RELB, but RELB overexpression improved tumor killing in the final two challenges (Fig. 3B). Flow cytometry analysis of TILs after the fourth tumor challenge revealed a dramatic increase in total CAR TIL cell numbers for two patient samples, while one patient sample failed to expand sufficiently for analysis (Fig. 3C-D, Supplemental Fig. 7A). These CAR TILs were mostly CD8+ (Fig. 3E-3F). CD8+ CAR TILs overexpressing RELB lacked PD-1 and LAG3 (Fig. 3F), while TIGIT was unaffected by RELB overexpression (Fig. 3F). Interestingly, comparing RELA to RELB overexpression demonstrated that RELA also conferred improved killing during later tumor challenges (Supplemental Fig. 7B), but RELA was unable to maintain CD8+ CAR TIL persistence (Supplemental Fig. 7C-D). It has been previously reported that CD4+ CAR-T cells have cytotoxic effector function^43^, which seems to be the main mechanism by which RELA-overexpressing TILs are acting given the lack of CD8+ CAR-TILs (Supplemental Fig. 7C-D).

**Fig. 3:**
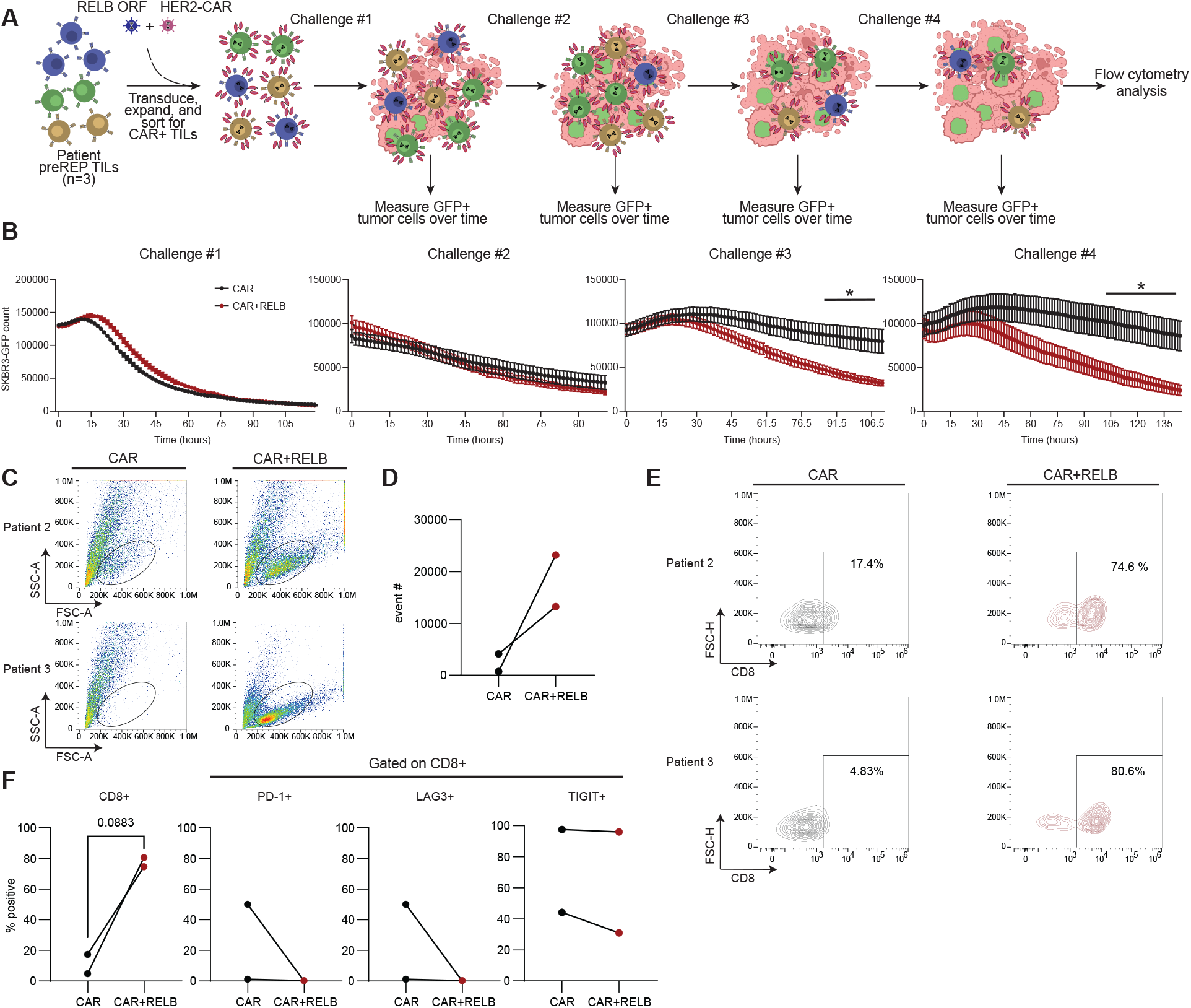
RELB improves CD8+ TIL persistence in tumor rechallenge model. (A) Experimental layout describing *in vitro* tumor rechallenge killing assay using a HER2-targetting CAR. (B) GFP+ count of SKBR3-GFP cells via live cell imaging over tumor multi-challenge assay (n = 3 distinct TIL patient samples, error bars represent standard error of the mean). * = p < 0.05 using two-way ANOVA analysis with Dunnett’s multiple comparisons test. (C) Flow cytometry analysis of side-scatter versus forward-scatter area of final rechallenge assay supernatant. (D) Total events count within side-scatter and forward-scatter gate. (E) CD8+ TIL events gated using fluorescence minus-one (FMO) controls. (F) Percent positive events for CD8+, CD8+ PD1+, CD8+ LAG3+, and CD8+ TIGIT+ based on flow gating on FMO controls. P value calculated for % CD8+ events with paired T-test.

### RELB improves TIL-mediated solid tumor killing *in vivo* and promotes TIL memory/costimulatory phenotypes

We then applied a similar tumor killing model *in vivo* by arming human TILs with a HER2-targeting CAR and injecting them into solid tumor-bearing mice (Fig. 4A). We used a 5×10^5^ CAR-TIL dose based on our previous dose-response studies with this model^35^. CAR-TILs from two patients were comprehensively analyzed by flow cytometry pre-injection at 21 days post-transduction (Fig. 4A, Supplemental Figure 8A), revealing that RELB overexpression in CD4+ and CD8+ TILs induced memory/stemness markers (IL7R for CD4s, CCR7 for CD8s, and TCF1 for both), the costimulatory receptor ICOS, and proliferation (Ki-67), but downregulated the exhaustion markers PD-1 and CD69. Interestingly, there was a significant increase in TGF-β+ CD4 TILs (Fig. 4B) which is a cytokine associated with maintenance of tissue-resident T cells^44-46^.

**Fig. 4:**
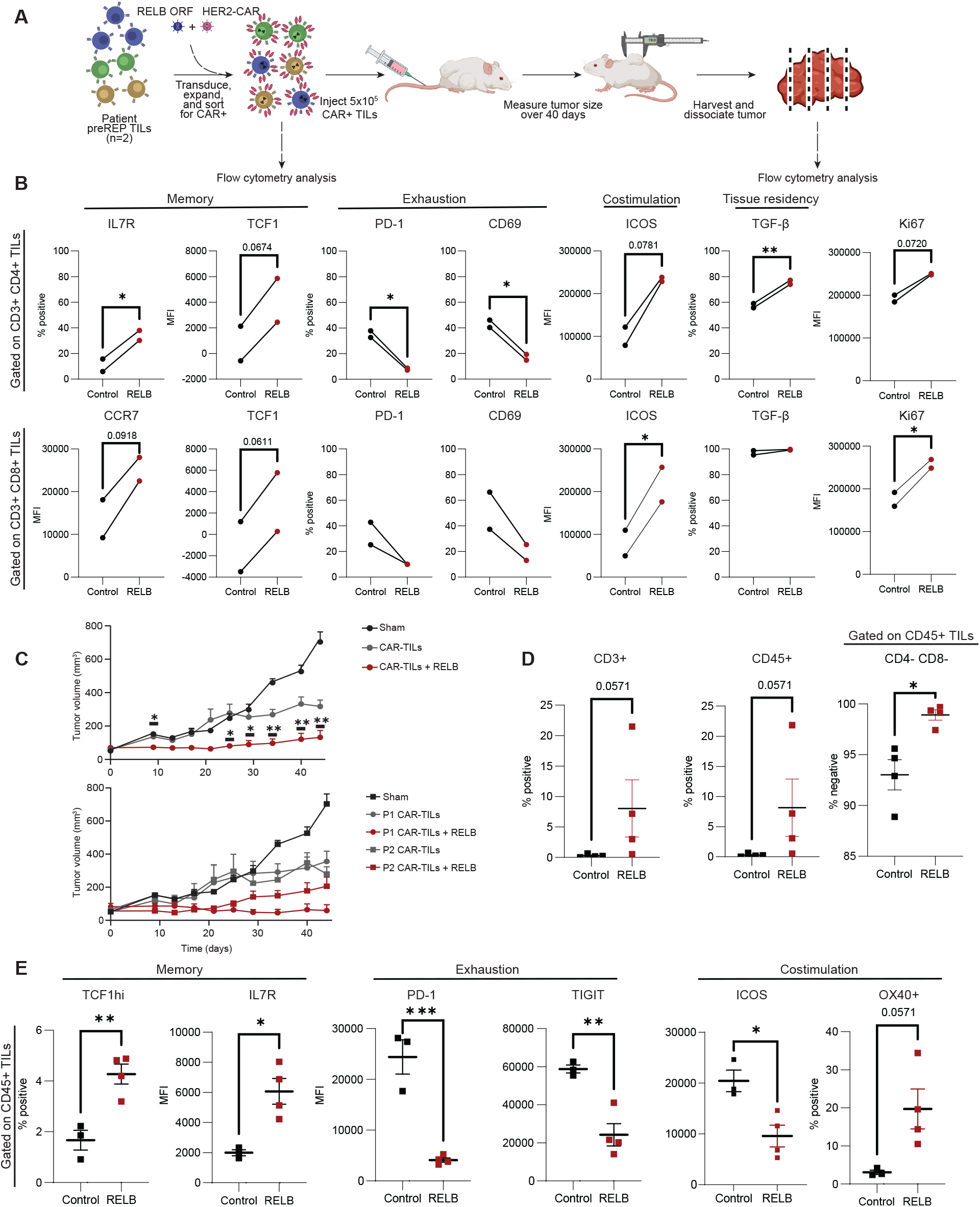
RELB improves TIL killing *in vivo* and promotes a more memory/costimulatory-like state in HER2 solid tumor model. (A) Experimental layout describing *in vivo* solid tumor challenge killing assay using a HER2-targetting CAR and flow cytometry analysis timeline pre-injection and post-tumor challenge (day 44 post CAR-TIL injection). (B) Flow cytometry phenotyping of pre-injection product for memory (IL7R, TCF1, CCR7, TCF1), exhaustion (PD-1, CD69), costimulation (ICOS), tissue-residency (TGF-β), and proliferation (Ki-67) markers. P values calculated using paired T-tests. * = p < 0.05 and ** = < 0.01. (C) Merged donor results of tumor volume (mm^3^) of untreated (n=4) and treated mice with 5×10^5 (n = 2 patient TIL samples, 4 mice per treatment). * = p < 0.05 between CAR-TIL + RELB and CAR-TIL conditions using a two-way ANOVA analysis with Dunnett’s multiple comparisons test (top panel). CAR-TIL control conditions were significantly different (p < 0.05) from untreated sham control at final three timepoints (top panel). Bottom panel shows unmerged results for both patient samples tested. (D) Percent positive CD3, CD45, and CD45+ CD4-CD8-CAR-TILs gated using FMO controls from final tumor harvest samples for one patient sample. P values calculated using Welch’s unpaired t test. * = < 0.05 p. value. (E) Percent positive CD45+ TCF1hi+, CD45+ IL7R+, CD45+ PD-1+, CD45+ TIGIT+, CD45+ ICOS+, CD45+ OX40+ CAR-TILs gated using FMO controls from final tumor harvest samples for one patient sample. P values calculated using Welch’s unpaired t test. * = p < 0.05, ** = p < 0.01, and *** = p < 0.001.

Notably, RELB overexpression in CAR-TILs led to a significant improvement in tumor control (Fig. 4C). Flow cytometry analysis of dissociated tumors 44 days post-treatment revealed a dramatic increase in CD3+ and CD45+ TILs (Fig. 4D, Supplemental Figure 8B) in tumors of mice receiving RELB expressing TILs. Surprisingly, nearly all CD45+ TILs were double-negative for CD4 and CD8 expression (Fig. 4D, Supplemental Figure 8B), possibly indicating a differentiation trajectory of the infused CD4+ and CD8+ CAR-TILs towards a memory-like CD4-CD8-double-negative (DN) state. Similar to our other observations, CD45+ DN TILs were maintained in a more memory-like state with a significant increase in memory markers (TCF1 and IL7R), significant downregulation of exhaustion markers (PD-1 and TIGIT), and a change in costimulatory proteins with significant downregulation of ICOS but a significant increase in OX40 expression (Fig. 4E, Supplemental Figure 8C). This DN state derived from peripheral CD4+ or CD8+ T cells has been characterized previously^47-49^, and given the high expression of OX40 these DN TILs are likely to have differentiated from CD4+ TILs^50,51^.

### RELB improves killing of patient-matched human tumor organoids

To assess patient-matched tumor killing with RELB-engineered TILs, we conducted an *in vitro* TIL and matched tumor organoid killing assay using flow cytometry to analyze T cell activation, exhaustion, cytotoxicity markers and tumor killing via LIVE/DEAD staining and Granzyme B levels (Fig. 5A). As we saw previously, there was a skew towards CD8+ TILs at the end of expansion with RELB overexpression compared to untransduced and control vector conditions (Supplemental Fig. 9A). Gating on CD3+ CD8+ TILs revealed a significant increase in 4-1BB and TIGIT for conditions overexpressing RELB while PD-1 expression was significantly decreased (Fig. 5B-C). Blocking MHC-I binding significantly decreased expression of 4-1BB and TIGIT for all patient samples (Fig. 5C), indicating that a significant fraction of CD8+ TILs were tumor-reactive via MHC-I binding and that RELB drove expansion of these tumor-reactive TILs.

**Fig. 5:**
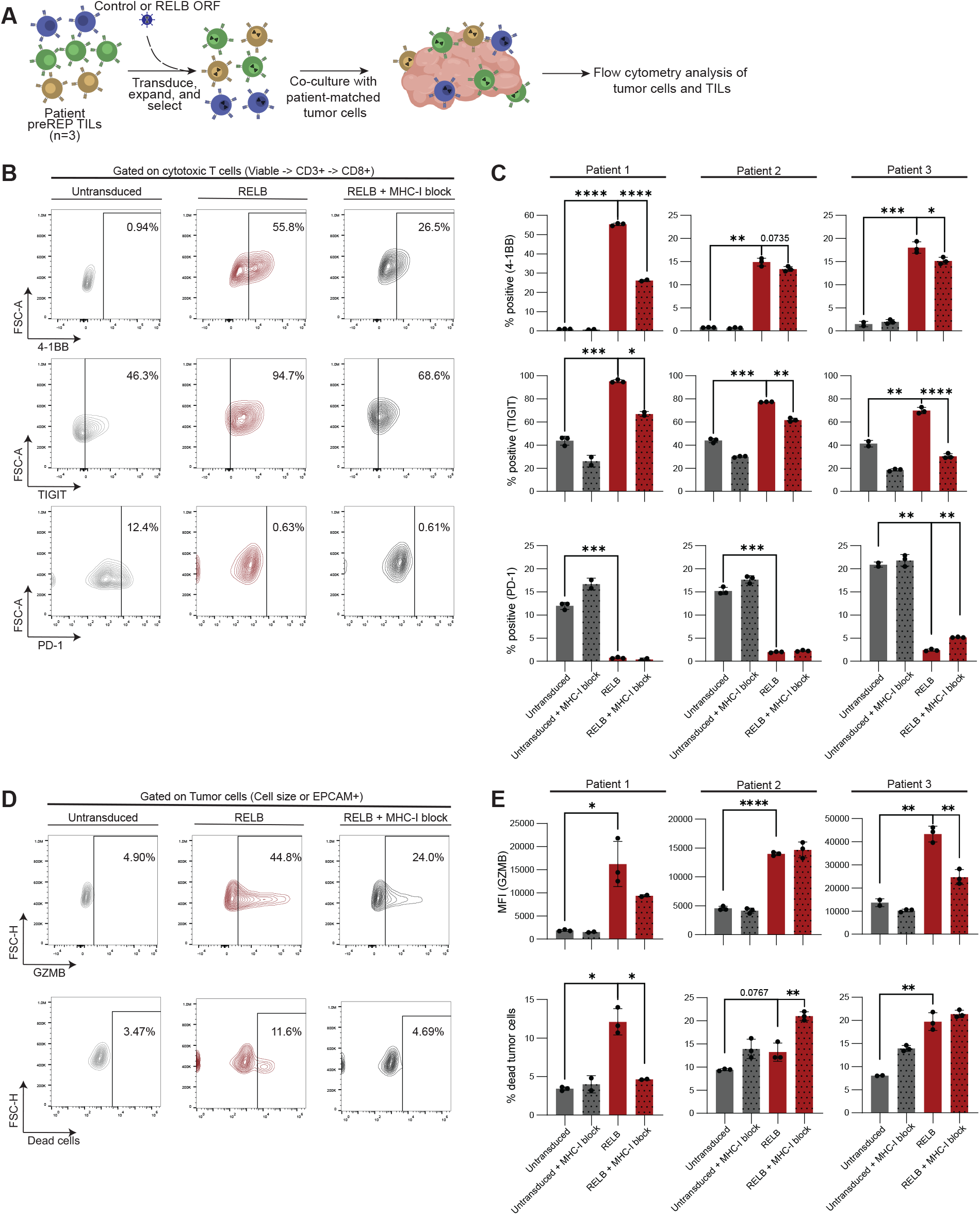
RELB improves TIL killing in matched tumor organoid co-culture assay *in vitro*. (A) Experimental layout describing *in vitro* TIL + matched tumor organoid co-culture killing assay conducted at 1:1 effector to target cell ratio. (B) Representative flow cytometry gating of viable CD3+ CD8+ TILs for T cell activation (4-1BB), and exhaustion (TIGIT, PD-1) markers. Gates were drawn based on FMO controls. (C) Summary data for all patient samples of CD3+ CD8+ TILs that were positive for 4-1BB, TIGIT, and PD-1. P values calculated using Welch’s unpaired t test. * = p < 0.05, ** = p < 0.01, *** = p < 0.001, **** = p < 0.0001. (D) Representative flow cytometry gating of tumor cells (based on either cell size – patient 1 or EPCAM+ signal – patients 2 and 3) for T-cell mediated cytotoxicity (Granzyme B) or viability (LIVE/DEAD). Gates were drawn based on FMO controls. (E) Summary data for all patient samples of tumor cells (gated on cell size or EPCAM+ signal) that were positive for Granzyme B or LIVE/DEAD signal. P values calculated using Welch’s unpaired t test. * = p < 0.05, ** = p < 0.01, *** = p < 0.001, **** = p < 0.0001.

Gating on NSCLC tumor cells via cell size and EPCAM+ signal revealed a significant increase in granzyme B levels within tumor cells for all patient samples in RELB-engineered TIL conditions compared to controls (Fig. 5D and 5E). Blocking MHC-I binding inhibited this increase in tumor-Granzyme B levels for two patients (Fig. 5E). Cytokine and effector protein secretion was also analyzed for two out of the three patients (due to sample limitations, cytokine expression was not measured for patient 1), demonstrating an increase in IFN-gamma, Granzyme A, Granzyme B, Perforin, and Granulysin for both patients in all conditions overexpressing RELB (Supplemental Fig. 9B-F). MHC-I block decreased secretion of all proteins for patient 3 (Supplemental Fig. 9B-F). All patient samples had an increased percentage of dead tumor cells with RELB-expressing TILs compared to controls (Fig. 5E). Interestingly, patient #2 had a significant increase in dead tumor cells when MHC-I binding was blocked, possibly indicating a different tumor-killing mechanism that is driving cytotoxicity in these TIL samples (Fig. 5E). Taken together, these results show that RELB overexpression improves the tumor-reactivity, tumor-killing, and polyfunctionality of NSCLC-derived TIL products.

## Discussion

High-throughput genetic screening of PBMC-derived human T cells has led to important discoveries of modulators of T cell phenotypes across many landmark studies^32-36,52^. However, the same technologies have not been extensively applied to human TILs and terminally exhausted T cells. We identified RELB as a potent positive regulator of exhausted TIL proliferation, which pushes terminally exhausted CD8+ CD103+ CD39+ TILs to a more memory and costimulatory-like state with improved persistence and tumor killing. The RELB open-reading frame size is only ∼1.7 kb, making it compatible with multiple delivery modalities that are readily adaptable to current TIL and CAR-T manufacturing contexts. Moreover, we showed that mRNA delivery of RELB via electroporation drives a similar phenotype that we see with lentiviral delivery.

To address concerns about uncontrolled TIL proliferation, we conducted two main experiments testing whether RELB overexpression led to either TCR-independent or cytokine-independent growth. RELB-transduced TILs that had not been activated with anti-CD3/CD28 showed no proliferation over 10 days *in vitro*. IL-2 deprivation also led to a halt in expansion of RELB-modified TILs and eventual death *in vitro*. Future studies will need to address any possible uncontrolled TIL proliferation *in vivo*.

TCR diversity of TILs was maintained throughout the RELB-induced *in vitro* expansion phase, excluding the possibility that only rare dominant clones were responding to RELB and overtaking the expanded cell population. However, we did not directly assess whether RELB overexpression was reprogramming specific TCR clones. To address this, future work could use the MANAFEST assay^53^ to identify tumor-specific TCR clones and then conduct *in vitro* patient-matched organoid co-culture experiments to track tumor-reactive TCR clones pre/post co-culture to determine if RELB overexpression is specifically enriching tumor-reactive clones.

While we extensively show that RELB leads to an improved cell product with TILs from NSCLC resections, this approach remains to be tested across different cancer types. Future studies could also establish patient-derived xenograft (PDX) models to assess RELB-expressing TIL reactivity against patient-matched tumors *in vivo*, to complement our current *in vivo* results with CAR-armed TILs. Additional studies may also compare or combine RELB overexpression with other next-generation T cell reprogramming approaches such as CISH knockout^54^, c-Jun overexpression^55^, REGNASE-1 knockout^56^, BATF overexpression^57^, and LTBR overexpression^34^. The robustness of TIL response to RELB across patient samples and assays supports its significance amongst the battery of tools necessary to address the difficult challenge of enabling ACT for solid tumor indications.

## Material and Methods

### Ethics statement

All animal experiments were conducted with strict adherence to the guidelines for the care and use of laboratory animals of the National Institutes of Health.

### Plasmids

All plasmids used for this work were obtained from Feng Zhang Lab through the Multiplexed Overexpression of Regulatory Factors (MORF) collection provided by Addgene (library – #192821, GFP plasmid – #145025, RELB plasmid – #142474). The Thy1.1 MORF control vector was generated by amplifying the Thy1.1 open-reading frame, double-digesting the GFP MORF plasmid (Addgene #145025) with NheI and SpeI restriction enzymes (NEB) and inserting the PCR product through Gibson assembly (NEB). A lentiviral vector encoding a HER2-targeting CAR with Myc epitope tag was used^58^. The ORF vectors with GFP reporter were generated by amplifying GFP, double digesting either the Thy1.1 MORF or RELB MORF plasmid (Addgene #142474) with KpnI and EcoRI restriction enzymes (NEB) and inserting the PCR product through Gibson assembly (NEB).

### Cell lines

HEK293T cells were maintained in Dulbecco’s modified Eagle medium (DMEM) GlutaMAX supplemented with 10% fetal bovine serum (FBS), 1 mM sodium pyruvate, 1× MEM non-essential amino acids, 10 mM HEPES, 100 U ml^-1^ penicillin and 100 μg ml^−1^ streptomycin. SKBR3s were maintained in McCoy’s supplemented with 10% FBS, 100 U ml^−1^ penicillin and 100 μg ml^−1^ streptomycin. HCC1954 cells were maintained in DMEM/F12 supplemented with 10% FBS, 100 U ml^−1^ penicillin and 100 μg ml^−1^ streptomycin.

### NSCLC patient tumor samples

All lung tumor samples were collected with consent under the IRB protocol Pro00102781. The samples were de-identified, placed into RPMI cell culture media and transported to the process laboratory on ice. The tumor tissue was divided in half. One half was used for tumor organoid generation, and the other half was used for TILs manufacturing. All samples were processed within 24 hours of collection.

### Isolation and culture of primary human NSCLC TILs

The weight of the solid tumor was recorded and the tumor cut to 2-3 mm^3^ pieces with a sterile scalpel. The fragments were placed into a G-Rex flask (Wilson-Wolf) at 2-20 g tissue per cm^2^ in preREP media (PrimeXV XSFM T-cell expansion media, FujiFilm or Excellerate T-cell media, Bio-Techne) supplemented with 5% human platelet lysate, (Compass Biotech), 1x Pen/Strep reagent (Gibco) and either 6000 IU/ml medical grade recombinant IL-2 (Proleukin, Iovance Biotherapeutics) or 4000 IU/ml GMP grade recombinant IL-2 (Bio-Techne). Cell culture lactate levels were monitored using a Lactate-Plus meter and lactate strips (Novus Biochemicals). Media was changed when the lactate concentration in the culture media reached 15mM or higher.

Expanded TILs were harvested on culture day 14 and yields determined with the automated cell counter Cellometer Auto2000 (Nexcelom/Revvity) using the manufacturer pre-installed program “immune cells, low RBC”. Harvested preREP TILs were pelleted at 300g for 10 min, re-suspended in CS10 (CryoStor, Biolife Solutions), cryopreserved using the CoolCell system (Corning Life Sciences) at −80°C and stored under liquid nitrogen (N2) vapor. At the end of preREP expansion TILs were frozen for future experiments.

All experiments described in this article begin with thaw of preREP TILs. preREP TILs were thawed and activated using 10 μL per mL of media of human T cell activator Transact (Miltenyi) at 1×10^6^ cells/mL. TILs were expanded in PRIME-XV T cell Expansion XSFM (Fujifilm Biosciences) supplemented with 5% human platelet lysate (Compass Biomedical), 100 U ml^−1^ penicillin and 100 μg ml^−1^ streptomycin. All media were supplemented with 1000 U ml^−1^ human IL-2 (Peprotech) unless otherwise stated.

### Lentivirus generation and transduction of TILs

For all TIL reprogramming experiments, a recently optimized transfection protocol was used^52^. Lentiviral supernatant was centrifuged at 600*g* for 10 min to remove cellular debris and concentrated to 50–100× the initial concentration using Lenti-X Concentrator (Takara Bio). T cells were transduced at 5–10% v/v of concentrated lentivirus at 24 h post-activation. For dual transduction experiments, TILs were co-transduced at 24 h post activation at 5–10% v/v of concentrated lentivirus.

### Flow cytometry and surface/cytoplasmic staining

For analyzing and sorting the ORF TF-ome FACS based screens, a BD Astrios cell sorter was used (Duke Cancer Institute Flow Cytometry Core). A SH800 FACS Cell Sorter (Sony Biotechnology) was used for cell sorting and analysis for all *in vitro* experiments except for the late mRNA delivery and matched organoid killing assay experiments. Instead, an NXT Attune flow cytometer (Thermo) was used for analysis of mRNA and matched killing experiments.

For antibody staining of all surface markers except CCR7, cells were collected, spun down at 300*g* for 5 min, resuspended in Cell Staining Buffer (Biolegend) with the appropriate antibody dilutions and incubated for 30 min at 4 °C on a rocker. Antibody staining of CCR7 was carried out for 30 min at 37 °C. Cells were then washed with flow buffer, spun down at 300*g* for 5 min and resuspended in flow buffer for cell sorting or analysis. Fluorescent minus one (FMO) controls were used to set appropriate gates for all flow panels.

For antibody staining of all intracellular markers, the Intraceullar Fixation & Permeabilization buffer (eBioscience, Invitrogen) set was used following manufacturer’s protocol. In brief, after surface staining, cells are spun down, resuspended in fixation buffer, and incubated at room temperature for 30 minutes. Afterwards cells were washed with permeabilization buffer, resuspended in permeabilization buffer with manufacturer-recommended antibody dilutions, and stained for 30 minutes at room temperature. Cells were washed twice with permeabilization buffer, resuspended in flow buffer and analyzed via flow cytometry.

All antibody details for every experiment can be found in Supplemental Table 4 and all relevant gating/analysis was conducted using Flow Jo V10.10.0.

### ORF TF-ome FACS-based proliferation screens

CD8+ TILs were isolated using a TIL CD8+ magnetic isolation kit (REAlease® CD8 (TIL) MicroBead Kit, human, Miltenyi) and CD8+ CD103+ CD39+ TILs were FACS sorted (n = 2 patient samples), activated with Transact (Miltenyi) according to manufacturer’s instructions, and transduced (n = 2 transduction replicates) with the MORF library at 8% v/v. Cells were expanded for 3 days, selected with puromycin at 1 µg/mL for 3 days, and then expanded for an additional 9 days. Cells were then re-activated at day 12 post-transduction with a 3:1 ratio of CD3/CD28 dynabeads (Invitrogen) to T cells, stained with CellTrace Violet (Invitrogen), and sorted for lower and upper 15% tails of CellTrace signal. All replicates were maintained and sorted at a minimum of 400x coverage. Cells were maintained at 1–2 × 10^6^ cells ml^−1^ for all expansion steps unless otherwise indicated.

### Genomic DNA isolation, gDNA PCR, and sequencing ORF libraries

Genomic DNA was isolated using Zymo’s QuickDNA Miniprep Plus kit. Genomic DNA was split across 100 μl PCR reactions (22 cycles at 98 °C for 10 s, 60 °C for 10 s, and 72 °C for 25 s) with NEBNext 2× Master Mix and up to 3 μg of genomic DNA per reaction. PCRs were pooled together for each sample and purified using double-sided SPRI bead selection at 0.5× and 1.0×. Libraries were run on a 1% agarose gel to confirm amplicon size and quantified using Qubit’s dsDNA High Sensitivity assay.

Libraries were diluted to 2 nM, pooled together at equal volumes, run on a High Sensitivity D1000 tape (Agilent) to confirm amplicon size, and sequenced using Illumina’s MiSeq Reagent Kit v2 (50 cycles). Primers used for the gDNA PCR are listed in Supplemental Table 4.

### Processing ORF barcode sequencing and ORF analysis for FACS-based proliferation screens

FASTQ files were aligned to custom indexes for each ORF barcode library (generated from the bowtie2-build function) using Bowtie 2^59^. Counts for each ORF barcode were extracted and used for further analysis in R. Individual ORF enrichment was determined using the DESeq2^60^ package to compare ORF abundance between groups for each screen. DESeq2 results are presented in Supplemental Table 1.

### ORF TIL proliferation validations

TILs from four NSCLC patients were thawed, activated with Transact (Miltenyi), and transduced 24 hours later with either the Thy1.1 ORF control vector or RELB ORF vector. Transduced TILs were selected with 1 ug/mL of Puro from day 2 to day 4 post transduction. On Day 4, 100k viable cells were sorted using LIVE/DEAD staining (Thermo) and plated. All conditions were then counted every 2-3 days using Thermo’s Countess 3 with Trypan blue staining for viability.

### ORF PBMC proliferation validations

CD8+ T cells were isolated from human PBMCs from three distinct donors (StemCell Tech) using a CD8+ negative magnetic selection kit (StemCell Tech), activated with anti-CD3/CD28 Dynabeads (Gibco), and transduced 24 hours later with either the Thy1.1 ORF control vector or RELB ORF vector. Transduced CD8+ T cells were selected with 1 μg/mL of Puro from day 2 to day 5 post transduction. On Day 6, T cells were stained using CellTrace Far Red and seeded at 10k cells/well into a poly-L-orthinine (Sigma-Aldrich) 96-well plate in cPRIME media (Fujifilm) supplemented with 100 U/mL IL-2. Images were captured with every hour for a total of 9 days of expansion using the Incucyte live-cell imaging system (Sartorius) with phase and red channels.

### TIL expansion in the absence of TCR activation

TILs from three NSCLC patients were thawed and transduced 24 hours later with either the Thy1.1 ORF + GFP control vector or RELB ORF + GFP vector. Four days after transduction, TILs were counted every two days for a total of 10 days. GFP expression was then analyzed via FACS.

### IL-2 deprived TIL expansion

TILs from four NSCLC patients were thawed, activated with Transact (Miltenyi), and transduced 24 hours later with either the Thy1.1 ORF control vector or RELB ORF vector. TILs were then expanded for three days and selected with 1 ug/mL of Puro for three days. TILs were then expanded for two days and seeded into a poly-L-orthinine (Sigma-Aldrich) 96-well plate in cPRIME media (Fujifilm) supplemented with/without 1000 U/mL IL-2 and 1000x dilution of Nuclight Red (Sartorius) to track proliferation with an Incucyte (Sartorius).

### mRNA delivery to TILs

For early RELB mRNA delivery, TILs from four NSCLC patients were thawed and activated with Transact (Miltenyi) in the presence of 1000 U/mL IL-2 at 1×10^6^ cells/mL. Three days post-activation, 1×10^6^ TILs were spun down, washed in PBS, and resuspended at 50×10^6^ cells/mL in 20 µL P3 buffer with supplement (Lonza Bioscience) containing either 1 μg of GFP mRNA (Trilink, unmodified) or RELB mRNA (pU modified). Cells were subsequently electroporated on a Lonza 4D Nucleofector X (Kit S cuvette, V4XP-3032) using the electroporation code EO115. Immediately after electroporation, 80 µl warm media was added to each electroporation well and cells were incubated for 15 min in a CO_2_ incubator at 37 °C. Cells were then transferred to a 12 well plate and brought to 1×10^6^ cells/mL in media supplemented 1000 U/mL IL-2. One day after electroporation, cells were stained in PBS with LIVE/DEAD Fixable Near-IR Dead Cell Stain Kit (Thermo, L10119) and sorted for live cells on a Sony sorter. GFP expression was measured to confirm mRNA delivery and translation (Supplemental Fig. 4A). Six days post-sort, TILs were stained with antibodies for IL7R, PD-1, LAG3, CD8, and TIGIT according to manufacturer’s recommendations and flow cytometry analyzed (antibody details in Supplemental Table 4).

For late RELB mRNA delivery, TILs from four NSCLC patients were thawed and activated with Transact (Miltenyi) in the presence of 1000 U/mL IL-2 at 1×10^6^ cells/mL. Fifteen days post-activation, TILs were re-activated with Transact (Miltenyi). Two days post-reactivation, 1×10^6^ TILs were spun down, washed in PBS, and resuspended at 50×10^6^ cells/mL in 20 µL P3 buffer with supplement (Lonza Bioscience) containing 1 μg of GFP mRNA (Trilink, M6 modified), or RELB mRNA (M6 modified) in a total of 20 µL. Cells were subsequently electroporated on a Lonza 4D Nucleofector X (Kit S cuvette, V4XP-3032) using the electroporation code EO115. Immediately after electroporation, 80 µl warm media was added to each electroporation well and cells were incubated for 15 min in a CO2 incubator at 37 °C. Cells were then transferred to a 12 well plate and brought to 1×10^6^ cells/mL in media supplemented 1000 U/mL IL-2. Two days post-electroporation, 1×10^4^ TILs of each condition were seeded in a poly-L-orthinine (Sigma-Aldrich) coated 96-well plate in cPRIME media (Fujifilm) supplemented with 1000 U/mL IL-2 and 1000x dilution of Nuclight Red (Sartorius) to track proliferation with an Incucyte (Sartorius). The plate was imaged every hour for four days. TILs were then harvested, stained according to manufacturer’s recommendations, and flow cytometry analyzed for Viability (LIVE/DEAD), CD3, CD4, CD8, CCR7, CD62L, IL7R, LAG-3, PD-1, 4-1BB, and 4-1BBL expression (antibody details in Supplemental Table 4). Fluorescent minus one (FMO) controls were used to set appropriate gates for all flow panels.

### TCR sequencing and data analysis

RNA was isolated using Norgen’s Total RNA Purification Plus Kit and Takara’s SMARTer Human TCR a/b Profiling Kit v1 was used to generate TCR-seq libraries. Library generation and purification methods were carried out according to manufacturer protocol. Initial data processing was performed on FASTQ files using MiXCR^61^ with the present for the SMARTer Human TCR a/b kit. The R package immunarch^62^ was used to analyze effective number of types for TCR diversity across samples to account for both number of clones and relative abundance.

### RNA sequencing and data analysis

RNA was isolated using Norgen’s Total RNA Purification Plus Kit and submitted to Azenta for standard RNA-seq with polyA selection. Reads were first trimmed using Trimmomatic^63^ v0.32 to remove adapters and then aligned to GRCh38 using STAR v2.4.1a aligner. Gene counts were obtained with featureCounts^64^ from the subread package (version 1.4.6-p4) using the comprehensive gene annotation in Gencode v22. Differential expression analysis was determined with DESeq2^60^ where gene counts are fitted into a negative binomial generalized linear model and a Wald test determines significant DEGs. DESeq2 results of RNA-seq analyses with RELB OE in all three TIL subtypes (CD4+, CD8+, CD8+ CD103+ CD39+) are presented in Supplemental Table 3. Relevant genes were plotted by calculating z-scores between all TIL subtypes from DESeq2 results using the R packages complexheatmap^65^ and ggplot2^66^. Enriched and depleted DEGs (p adj. <0.05) were used as input into EnrichR’s GO Biological Processes 2021 database^67^ for functional annotation; results are provided in Supplemental Table 3. For GSEA^68^, RELB vs Thy1.1 DESeq2 results were shrunk and used as input to msigdbr^69^ and clusterProfiler^70^, then tested against the C7 RNA-seq databases; results are provided in Supplemental Table 3. Relevant enriched/depleted gene pathways and RNA-seq datasets were plotted using R package ggplot2.

### *In vitro* CAR-TIL tumor rechallenge assay

TILs were thawed, activated with Transact (Miltenyi), and co-transduced 24 hours later with HER2-CAR lentivirus and either Thy1.1 ORF, RELB ORF, RELA ORF, or no virus (CAR only conditions). Co-transduced TILs were then puromycin selected for 3 days and expanded for an additional four days. CAR+ TILs were isolated by staining with a MYC:PE antibody (Supplemental Table 4) and FACS sorted. SKBR3 cells tagged with nuclear GFP were seeded on a 48-well plate in TIL media for Incucyte imaging the same day as CAR+ TIL sorting. Sorted CAR+ TILs were maintained for an additional day and seeded on Day 11 post-cotransduction at 1:4 TIL to cancer cell ratios. After nine days of co-culture, the entire supernatant was transferred to a fresh plate with SKBR3 cells and co-cultured for an additional four days. This was repeated two additional times. Supernatant was then harvested six days after the final challenge for flow cytometry analysis of CAR+ TILs with CD8, PD-1, LAG-3, and TIGIT antibodies (antibody details in Supplemental Table 4).

### Mice

All experiments involving animals were conducted with strict adherence to the guidelines for the care and use of laboratory animals of the National Institutes of Health. All experiments were approved by the Institutional Animal Care and Use Committee at Duke University (protocol number A130-22-07). Six-to 8-week-old female immunodeficient NOD/SCID gamma (NSG) mice were obtained from Jackson Laboratory and then housed in 12-h light/dark cycles, at an ambient temperature (21 ± 3 °C) with relative humidity (50 ± 20%) and handled in pathogen-free conditions. Mice were euthanized before reaching a tumor volume of 2,000 mm^3^, the upper threshold defined by the Duke Institutional Animal Care and Use Committee.

### *In vivo* tumor model

A total of 2.5 × 10^6^ HER2+ HCC1954 cells were implanted orthotopically into the fourth mammary fat pad of NSG mice in 100 μl 50:50 (v:v) PBS (Gibco):Matrigel (Corning #356237). Twenty-one days after tumor implantation, immediately before CAR TIL injections, mice were randomized into groups and tumors were measured. Tumor volumes were calculated on the basis of caliper measurements using the following formula: volume = ½(Length × Width^2^). Tumor measurements were performed blinded to treatment group. TILs were expanded for 21 days post-cotransduction before treatment. TILs were stained with a MYC:PE antibody (Supplemental Table 4) to detect CAR expression. Transduced CAR+ TILs were sorted using a BD Astrios (Duke Cancer Institute Flow Cytometry Core) by gating on GFP+ for ORF expression and MYC+ signal for CAR expression. CAR TILs were resuspended at 2.5 × 106 cells ml^−1^ in 1× PBS (Gibco) for 200-μl intravenous (i.v.) injections of 5 × 10^5^ HER2 CAR+ TILs.

### Flow cytometry analysis of input and tumor-infiltrating CAR TILs

Mice bearing HCC1954 tumors were euthanized on day 41 post CAR TIL delivery under deep isoflurane anesthesia via exsanguination. Tumors were resected, minced and incubated in RPMI-1640 medium (Gibco) for 45 min in 100 µg/ml Liberase-TM (Sigma-Aldrich) and 10 µg/ml DNase I (Roche) at 37 °C. Single-cell suspensions were filtered through a 70-mm cell strainer (Olympus Plastics), washed in PBS, stained with Zombie NIR (1:250, BioLegend), washed in FACS buffer (2% FBS (Sigma) + PBS), and treated with 1:50 mouse and human Tru-stain Fc block (both BioLegend). Cells were then stained for cell surface markers followed by intracellular staining using the Transcription Factor Staining Buffer Set (Invitrogen) per the manufacturer’s instructions. All data were collected on a Fortessa X 20 (Duke Cancer Institute Flow Cytometry Core; Supplemental Table 4 - Panel 1) or a Cytek Aurora (Supplemental Table 4 - Panel 2). De-identified leukopak-derived (Stemcell Tech) PBMCs or antibody capture beads (Beckman-Coulter) were used for spectral unmixing as indicated in Supplemental Table 4. Samples with less than 500 recovered T cells were excluded from T cell phenotyping analyses. Blood/tumor from sham-infused mice and FMO controls were used to guide gating for CAR T cells and to confirm appropriate compensation/unmixing, respectively.

### NSCLC organoid generation

Previously described tumor tissue samples were dissociated mechanically and enzymatically in 2 mg/mL of collagenase (Sigma), 600 U/mL of DNase (Sigma), and 1 protease inhibitor tablet (Roche) in serum-free RPMI media (Sigma-Aldrich) for 1 to 3 hours while rocking. Undigested tumor fragments were pressed through 70-µm mesh and added to a digested single-cell suspension. Cells were pelleted by centrifugation, resuspended in PBS, and cell viability was determined by trypan blue.

Single-cell suspension of dissociated tumor tissue was used for the 3D organoid culture in accordance with published methods^71,72^. Cells were resuspended with Matrigel (Corning) and 100 μl Matrigel plus cells was placed in the middle of a well of 24-well tissue culture plate. Matrigel droplets were solidified at 37°C for 10-20 min. Lung tumor organoid specific medium was prepared by supplementing Advanced DMEM/F12 medium with GlutaMax, Hepes, Penicillin/Streptomycin (all from Gibco) and Primocin (Invivogen), 500 ng/ml of R-Spondin-1, 25 ng/ml of FGF-7, 100 ng/ml of FGF-10, 100 ng/ml of Noggin (all from Peprotech), 500 nM of A83-01, 5 µM of Y-27632 (both Tocris), 1XB27 supplement (Gibco), 500 nM of SB202190, 1.25 mM of N-Acetylcysteine and 5 mM of Nicotinamide (all from Sigma Aldrich). Fresh supplemented medium was added to the solidified domes, and plates were transferred to 5% CO2 incubator for organoid generation. Tumor organoid lines were established within 2 months of culture and cryopreserved in Recovery Cell Culture Freezing Medium (Thermo Fisher) as master stock for a biobank.

### *In vitro* matched tumor organoid killing assay

TILs from three NSCLC patients were thawed, activated with Transact (Miltenyi), and transduced 24 hours later with either Thy1.1 ORF control vector or RELB ORF vector. TILs were then expanded for two (patient 1) or three days (patients 2 and 3) and selected with 1 ug/mL of Puro for two (patient 1) or three days (patients 2 and 3). TILs were then expanded for an additional seven days. Matched tumor organoids were dissociated using TryPLE and 1×10^5^ were seeded in a 24-well plate in airway organoid media one day before co-culture start. MHC-I binding was blocked incubating seeded organoid cells with 2-5 µg/mL anti HLA-A,B,C antibody (Biolegend) for 30 minutes at 37 C before co-culture start. TILs were seeded into organoid plates at 1:1 TIL to organoid cell ratio in triplicate and co-cultured for three days. Both TILs from the supernatant and adherent tumor cells were harvested by using Accutase (StemCell Technologies). For two out of the three patient samples, a fraction of the supernatant was harvested for cytokine expression analysis using the Legendplex NK/CD8 bead-based assay (Biolegend). TILs and tumor cells were then flow cytometry analyzed for viability (LIVE/DEAD), CD3, CD4, CD8, EPCAM (for patients 2 and 3 only), 4-1BB, CD69, LAG-3, PD-1, TIGIT, IFN-γ, and Granzyme B (Supplemental Table 4).

### Statistics and reproducibility

All experiments utilized unique biological replicates due to the limited availability of primary human NSCLC TIL material. All statistical analysis methods are indicated in the figure legends (NS, not significant; \**P* < 0.05, \*\**P* < 0.01, \*\*\**P* < 0.001, \*\*\*\**P* < 0.0001).

Statistical analyses and data visualizations were performed in Graphpad Prism v.10.4.1 and R v4.2.1. All experiments have been replicated with at least two biological replicates. For in vivo studies, mice were randomly assigned into treatment groups. In this study, no statistical method was used to predetermine sample size, no data were excluded from the analyses, experiments were not randomized, and investigators were not blinded to allocation during experiments and outcome assessment.

## Data availability

All data associated with this study are present in the manuscript, supplementary information files, or have been deposited in the Gene Expression Omnibus (GEO) under accession numbers GSE303432 (proliferation screen), GSE303438 (TCR-seq), GSE303442 (RNA-seq).

## Acknowledgements

We thank all Gersbach and Antonia laboratory members for all the invaluable help provided for this project. We thank W. Wong for generously providing the HER2 CAR plasmid. Schematics in Fig. 3A, 4A, and 5A were created using Biorender. This work was supported by the Duke-Coulter Translational Partnership, NIH grants R01CA289574, UM1HG012053, RM1HG011123, and T32GM142605, and Yosemite Management, LLC. Supporting centers include the Duke Cancer Institute Flow Cytometry Core, Sequencing and Genomic Technologies Core, and the Duke Center for Genomic and Computational Biology.

## Conflict of interest statement

C.A.G. is a co-founder of Tune Therapeutics, Sollus Therapeutics, and Locus Biosciences and is an advisor to Tune Therapeutics, Sarepta Therapeutics, and Pappas Capital, LLC. S.A. is an advisor to Immutep and a co-founder of Cellepus Therapeutics. C.D.M.A., S.R.J., and C.A.G. are inventors on patents or patent applications related to CRISPR editing technologies, methods for high-throughput screening, and engineering of T cell therapies.

## Notes

https://www.ncbi.nlm.nih.gov/geo/query/acc.cgi?acc=GSE303432

https://www.ncbi.nlm.nih.gov/geo/query/acc.cgi?acc=GSE303438

https://www.ncbi.nlm.nih.gov/geo/query/acc.cgi?acc=GSE303442

