## Supplemental Figures for "RELB Reprograms Exhausted Tumor-Infiltrating Lymphocytes for Improved Adoptive Cell Therapy"

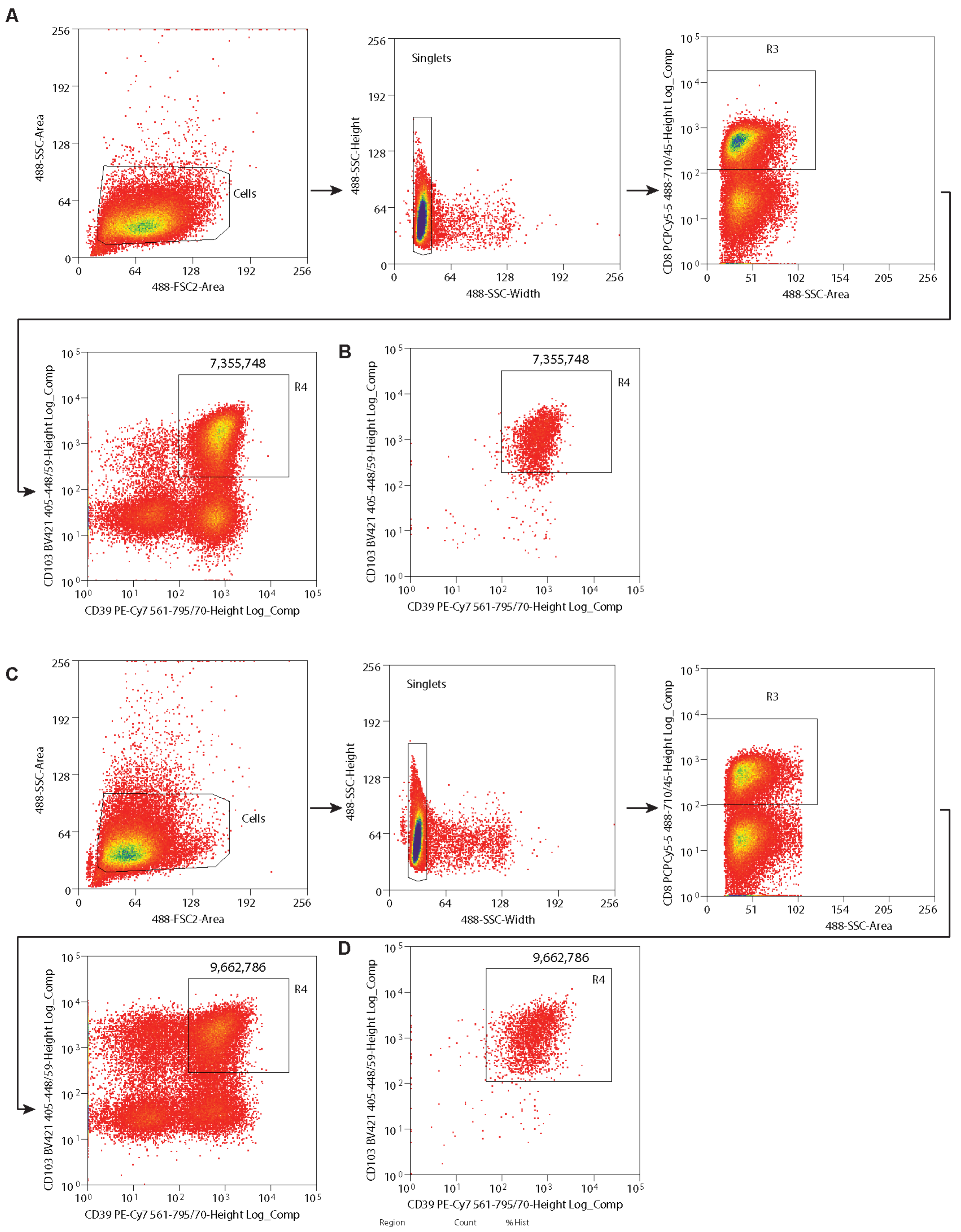

A
Cells
Singlets
R3
7,355,748
R4
B
C
9,662,786
D
488-SSC-Area
488-FSC2-Area
488-SSC-Height
488-SSC-Width
CD8 PCPCy5-5 488-710/45-Height Log_Comp
CD103 BV421 405-448/59-Height Log_Comp
CD39 PE-Cy7 561-795/70-Height Log_Comp
Region
Count
%Hist

**Supplemental fig. 1: Representative gating strategy for CD8+ CD103+ CD39+ TIL sorting.** (A) FACS gating strategy for CD8+ CD103+ CD39+ TILs from patient sample #1, arrows indicate what direction gates fed into. (B) Post-sort FACS analysis of CD103+ CD39+ TILs from patient sample #1. (C) FACS gating strategy for CD8+ CD103+ CD39+ TILs from patient sample #2, arrows indicate what direction gates fed into. (D) Post-sort FACS analysis of CD103+ CD39+ TILs from patient sample #2.

A

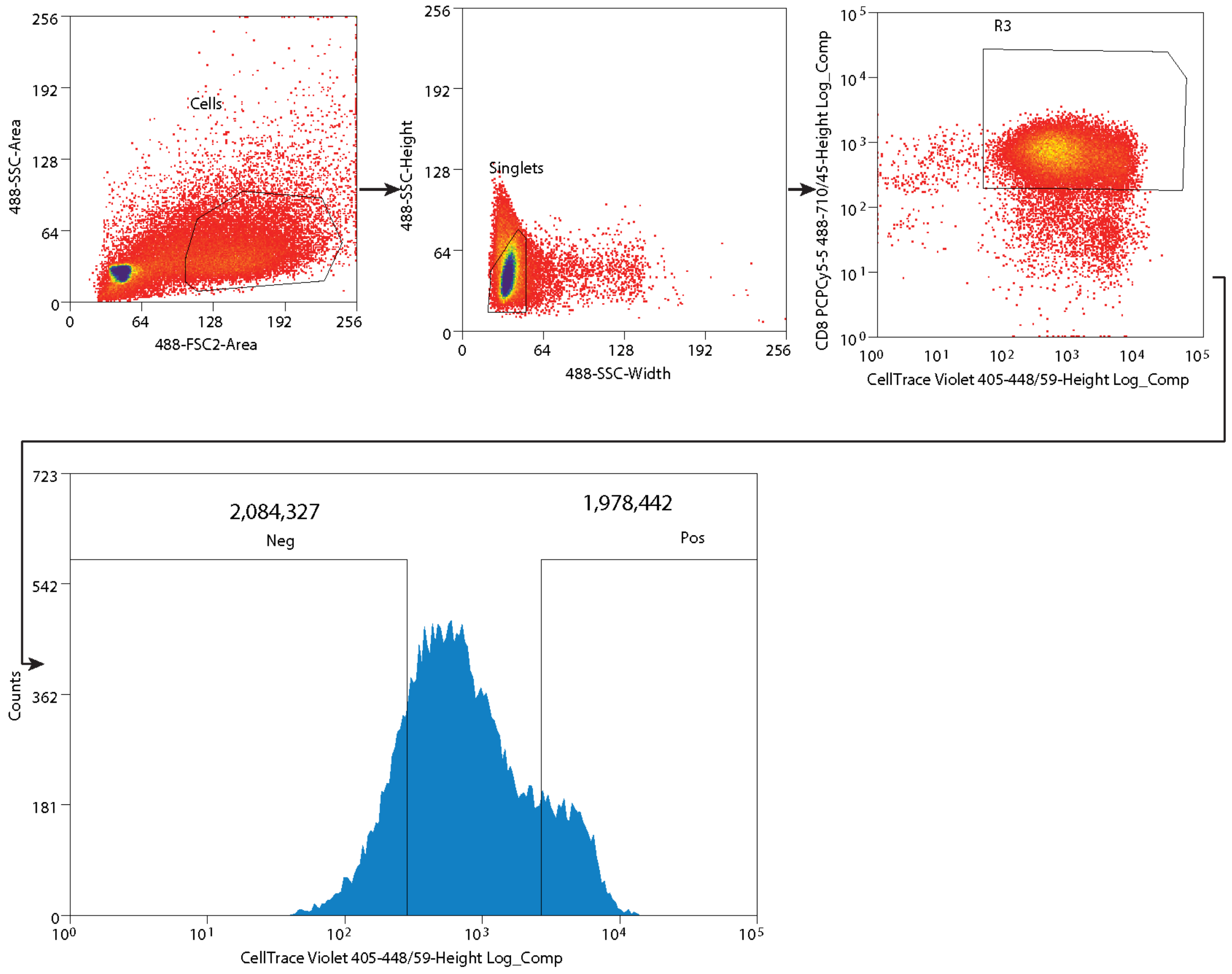

488-SSC-Area
Cells
488-FSC2-Area
488-SSC-Height
Singlets
488-SSC-Width
CD8 PCPCy5-5 488-710/45-Height Log_Comp
R3
CellTrace Violet 405-448/59-Height Log_Comp
Counts
2,084,327
Neg
1,978,442
Pos
CellTrace Violet 405-448/59-Height Log_Comp


B

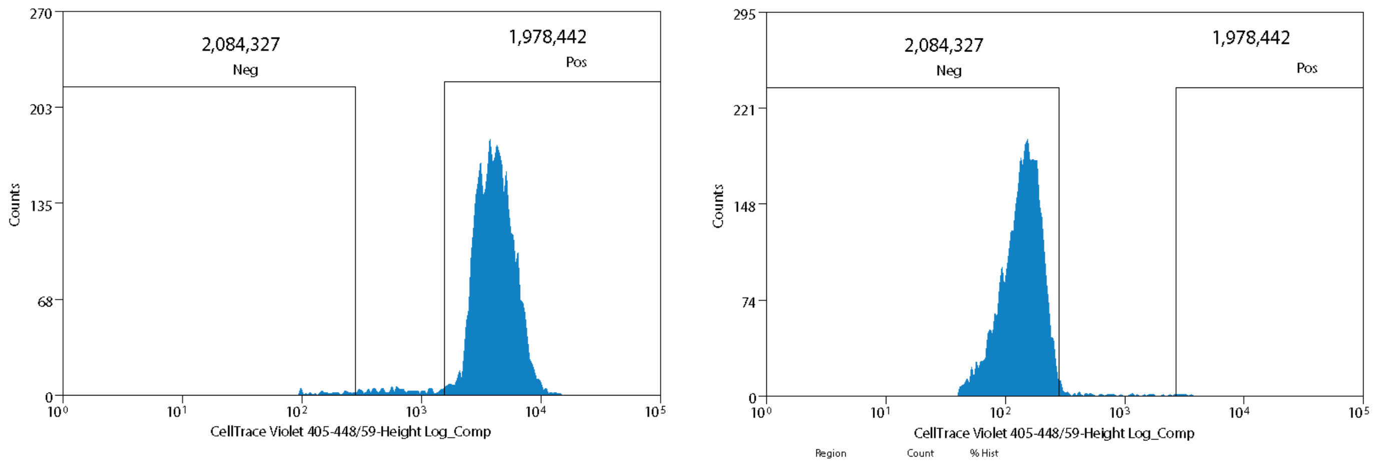

Counts
2,084,327
Neg
1,978,442
Pos
CellTrace Violet 405-448/59-Height Log_Comp
Counts
2,084,327
Neg
1,978,442
Pos
CellTrace Violet 405-448/59-Height Log_Comp
Region
Count
% Hist

**Supplemental fig. 2: Representative gating strategy for CellTrace-based TIL proliferation screen sort.** (A) FACS gating strategy for CellTrace high (top 15% CellTrace signal) versus low (bottom 15% CellTrace signal) TILs, arrows indicate what direction gates fed into. (B) Representative post-sort FACS analysis of CellTrace high TILs and CellTrace low TILs.

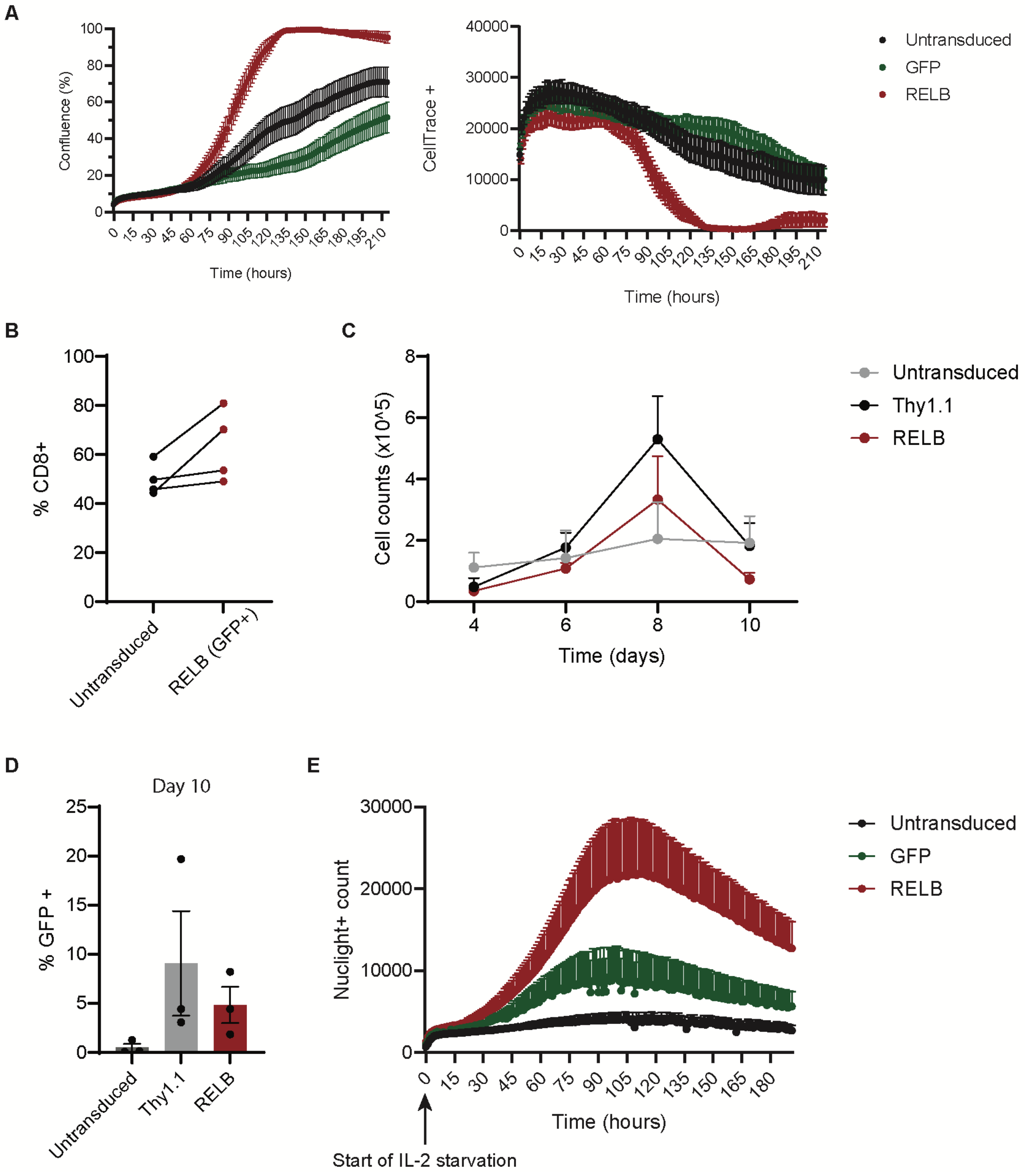

**Supplemental fig. 3: PBMC-derived CD8+ proliferation, GFP reporter proliferation, no TCR stimulation and IL-2 depravation proliferation experiments.** (A) Mean Incucyte confluence percentage and Celltrace+ counts for three distinct PBMC-derived CD8+ T cells (n = 3 donors, error bars represent standard error of the mean) over 10 day period. (B) Flow cytometry analysis of percent CD8+ TILs in untransduced and GFP+ RELB TILs 16 days post-transduction for four separate TIL patient samples. (C) Mean cell counts for four distinct TIL patient samples (error bars represent standard error of the mean) during 10 day *in vitro* expansion with no TCR activation. (D) Flow cytometry analysis of GFP expression on final day of expansion for no TCR activation experiment. (E) Mean Incucyte Nuclight red+ counts for four distinct TIL patient samples (error bars represent standard error of the mean) over 8 day period in culture media with no IL-2 supplementation.

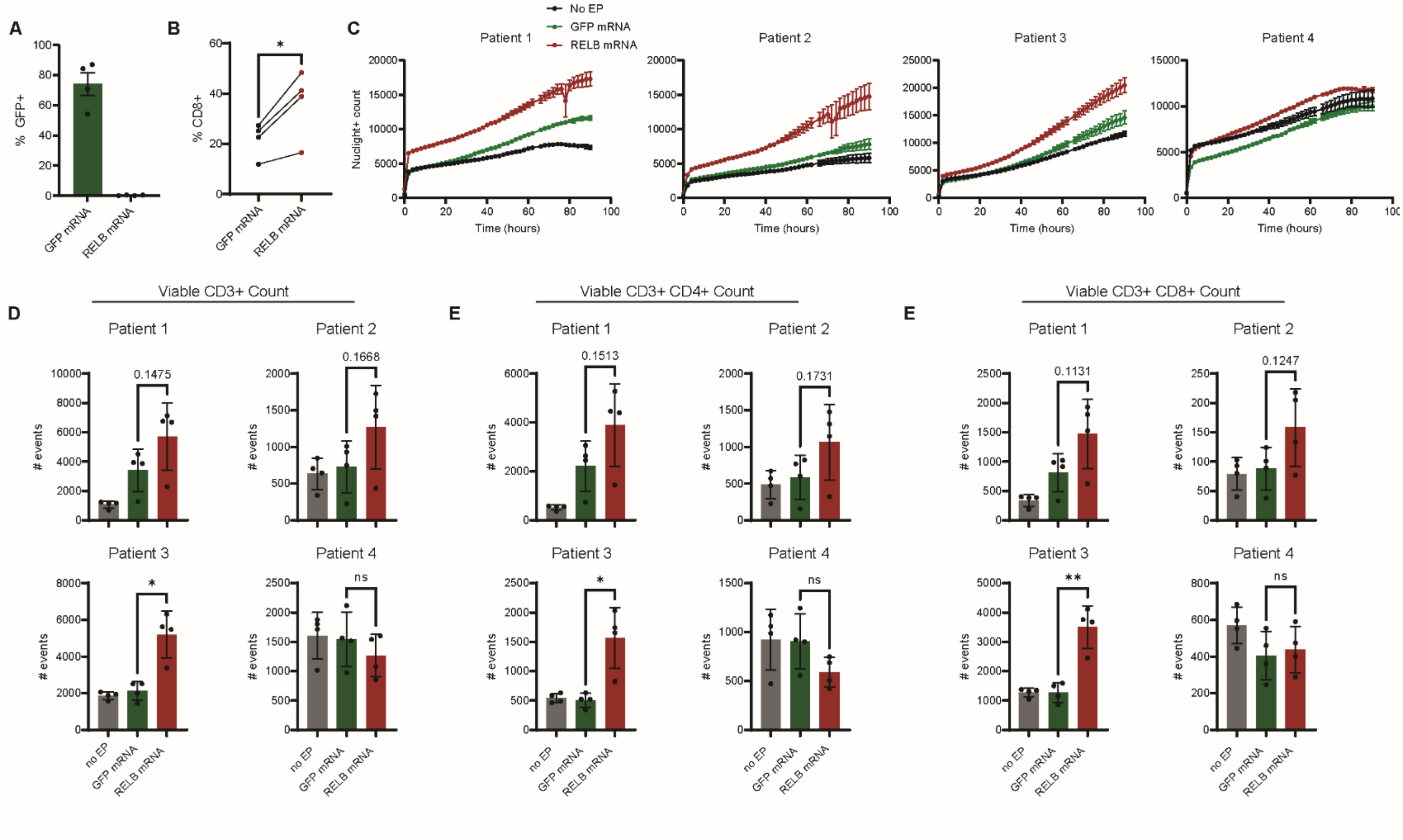


**Supplemental fig. 4: Transient delivery of RELB mRNA to human TILs.** (A) GFP expression was measured by flow cytometry to confirm mRNA delivery and translation one day post mRNA electroporation. RELB mRNA did not contain GFP reporter. (B) Percent positive CD8 TILs seven days post electroporation, ten days post initial activation. Gating was based on unstained control. * = <0.05 p. value using paired T-test. (C) Mean Incucyte Nuclight red+ counts for four TIL patient samples (n= 4 technical replicates) over 4 days of culture; experiment was plated two days post electroporation, seventeen days post initial activation. Error bars represent standard deviation. (D) Viable CD3+, (E) CD3+ CD8+, or (F) CD3+ CD4+ TIL events for all samples at the end of proliferation experiment. Error bars represent standard deviation. P. values were calculated using Welch’s unpaired T test. * = p. val < 0.05 and ** = < 0.01.

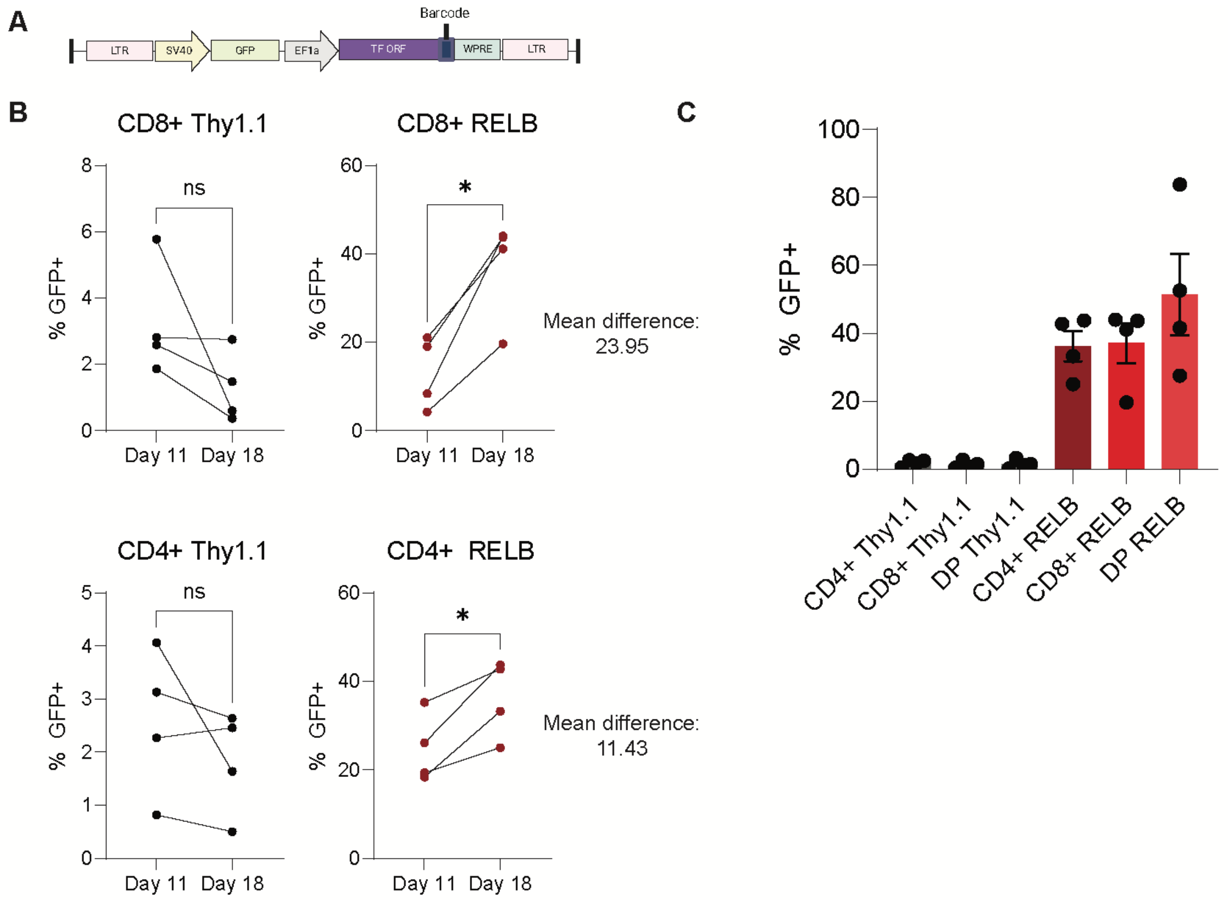


**Supplemental fig. 5: Proliferation of different human TIL subtypes.** (A) Lentiviral vector schematic for TF ORF vector co-expressing GFP as a reporter. (B) Percent GFP+ TILs for four patient samples as measured by flow cytometry in either CD4+ sorted or CD8+ sorted TILs for days 11 and 18 post-transduction of either Thy1.1 or RELB lentiviral vectors with GFP as a reporter. P. values and mean differences were calculated using paired T tests; * = <0.05 p. value. (C) Mean percent GFP+ TILs on final day of expansion (Day 18 post-transduction) for all sorted TIL subsets (error bars represent standard error of the mean).

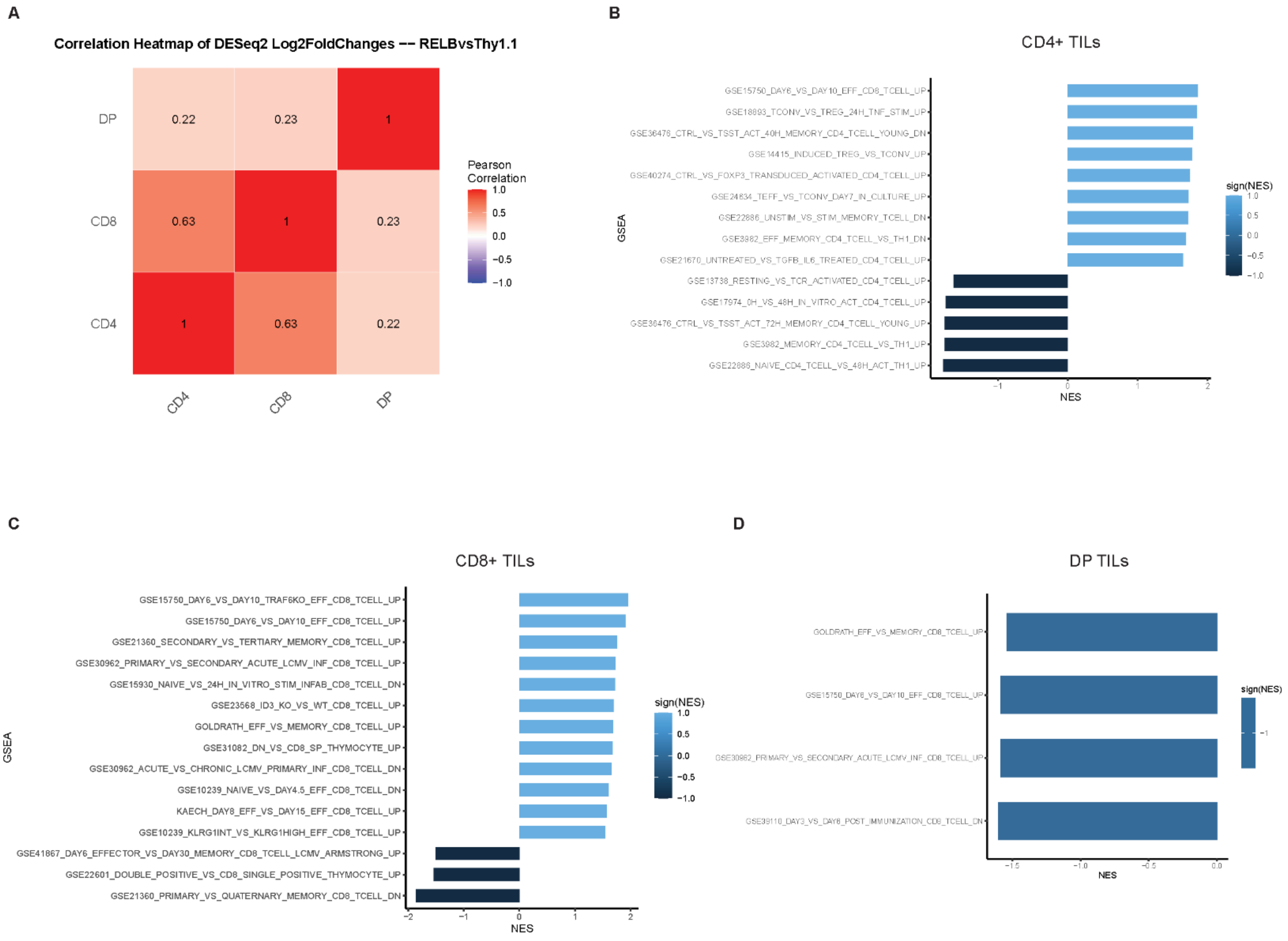


**Supplemental fig. 6: RNA-seq dataset correlation and GSEA enrichment analysis.** (A) Heatmap of Pearson correlation values between different sorted TIL subtypes for fold changes from RELB vs Thy1.1 ORF DESeq2 comparison. (B-D) Detailed names of RNA-seq datasets from GSEA of enriched and depleted gene sets in (B) CD4+, (C) CD8+, and (D) DP TILs (adj. p. val < 0.05). Positive NES corresponds to enrichment of pathway in RELB-overexpressing conditions while a negative NES corresponds to depletion.

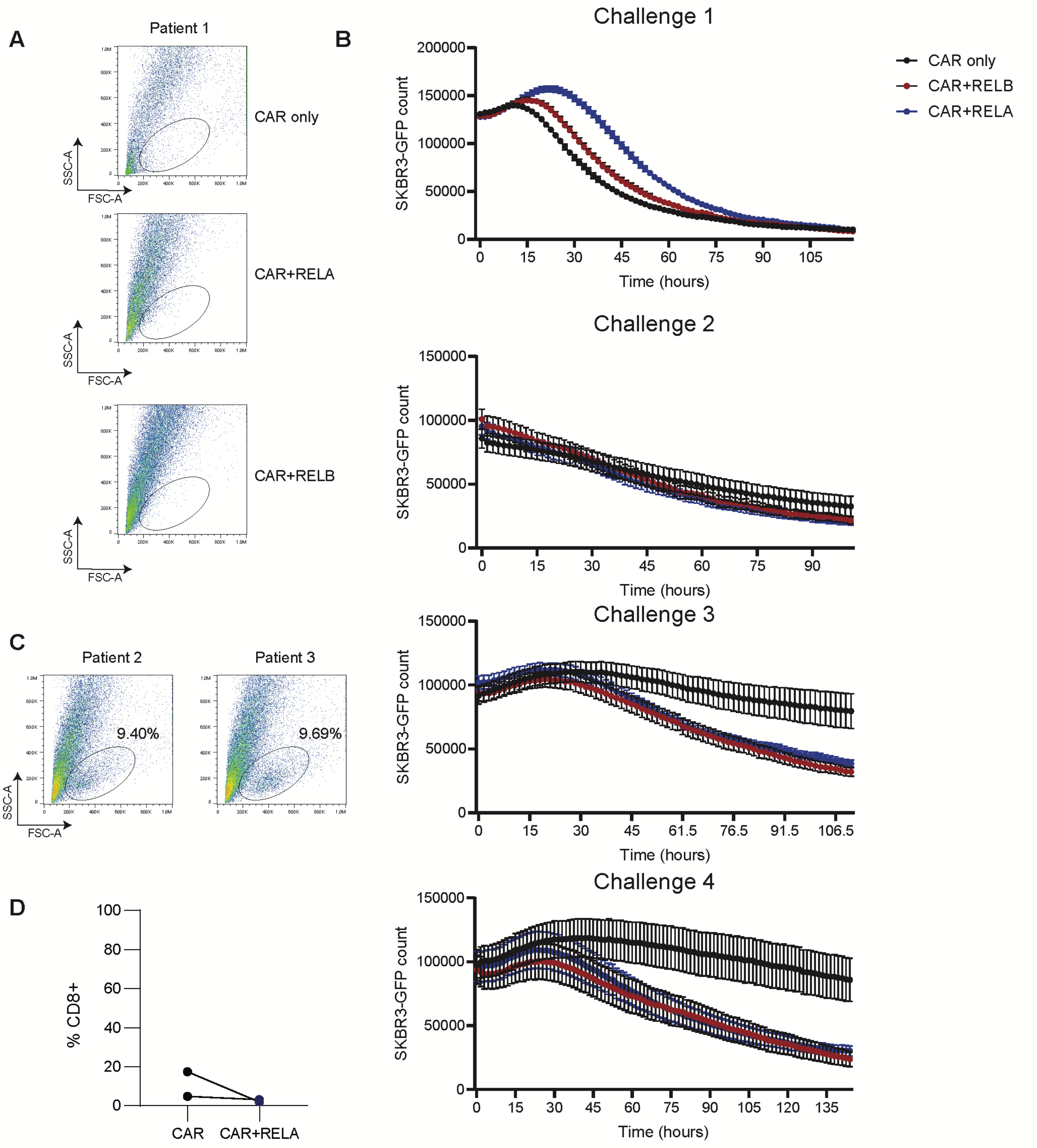


**Supplemental fig.7: Tumor rechallenge assay for RELA overexpression in TILs.** (A) Flow cytometry analysis of side-scatter versus forward-scatter area of final rechallenge supernatant for patient 1. (B) GFP+ count of SKBR3s-GFP with Incucyte live cell imaging system over tumor multi tumor challenge assay. (C) Flow cytometry analysis of side-scatter versus forward-scatter area of final rechallenge supernatant for RELA conditions (patients 2 and 3). (D) Percent CD8+ TIL events gated using fluorescent minus-one (FMO) controls.

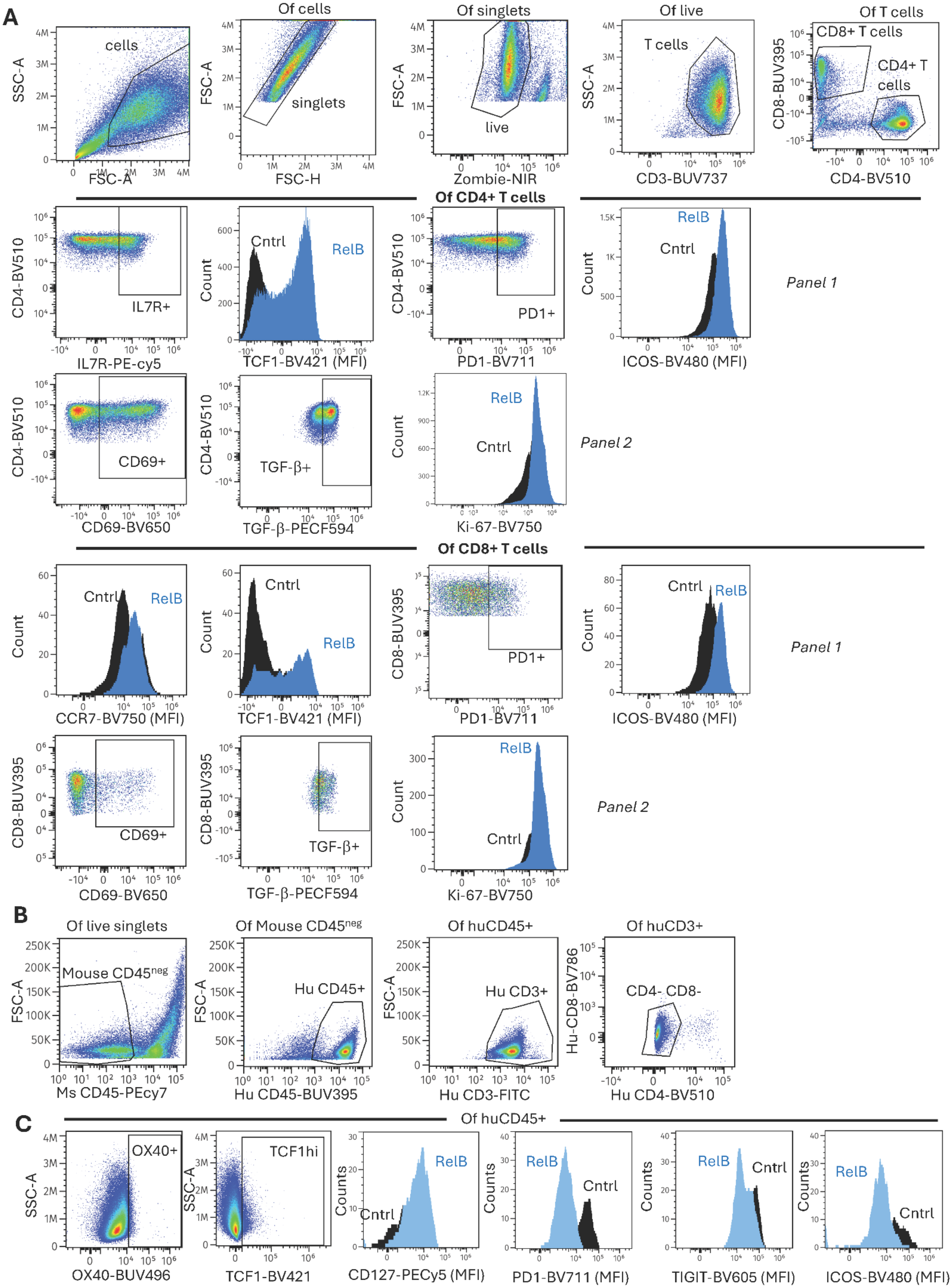
A
Of cells
Of singlets
Of live
Of T cells
cells
singlets
live
T cells
CD8+ T cells
CD4+ T cells
SSC-A
FSC-A
FSC-H
Zombie-NIR
CD3-BUV737
CD8-BUV395
CD4-BV510
Of CD4+ T cells
IL7R+
IL7R-PE-cy5
Cntrl
RelB
TCF1-BV421 (MFI)
PD1+
PD1-BV711
ICOS-BV480 (MFI)
Count
Panel 1
CD69+
CD69-BV650
TGF-β+
TGF-β-PECF594
Ki-67-BV750
Panel 2
Of CD8+ T cells
CCR7-BV750 (MFI)
B
Of live singlets
Of Mouse CD45neg
Of huCD45+
Of huCD3+
Mouse CD45neg
Hu CD45+
Hu CD3+
CD4- CD8-
Ms CD45-PEcy7
Hu CD45-BUV395
Hu CD3-FITC
Hu CD4-BV510
Hu-CD8-BV786
C
Of huCD45+
OX40+
TCF1hi
OX40-BUV496
TCF1-BV421
Counts
CD127-PECy5 (MFI)
PD1-BV711 (MFI)
TIGIT-BV605 (MFI)
ICOS-BV480 (MFI)

**Supplemental fig. 8: Representative gating strategy for input and in vivo TIL flow cytometry shown in Fig 4.** (A) Analysis of input TILs gated on CD4+ or CD8+ T cell subsets. (B) For identification of TILs, live singlets were gated as shown in (A) followed by removal of mouse CD45+ cells and gating on human CD45+ CD3+ cells. (C) Representative gating for phenotypes of TILs recovered in vivo. MFI= median fluorescence intensity.

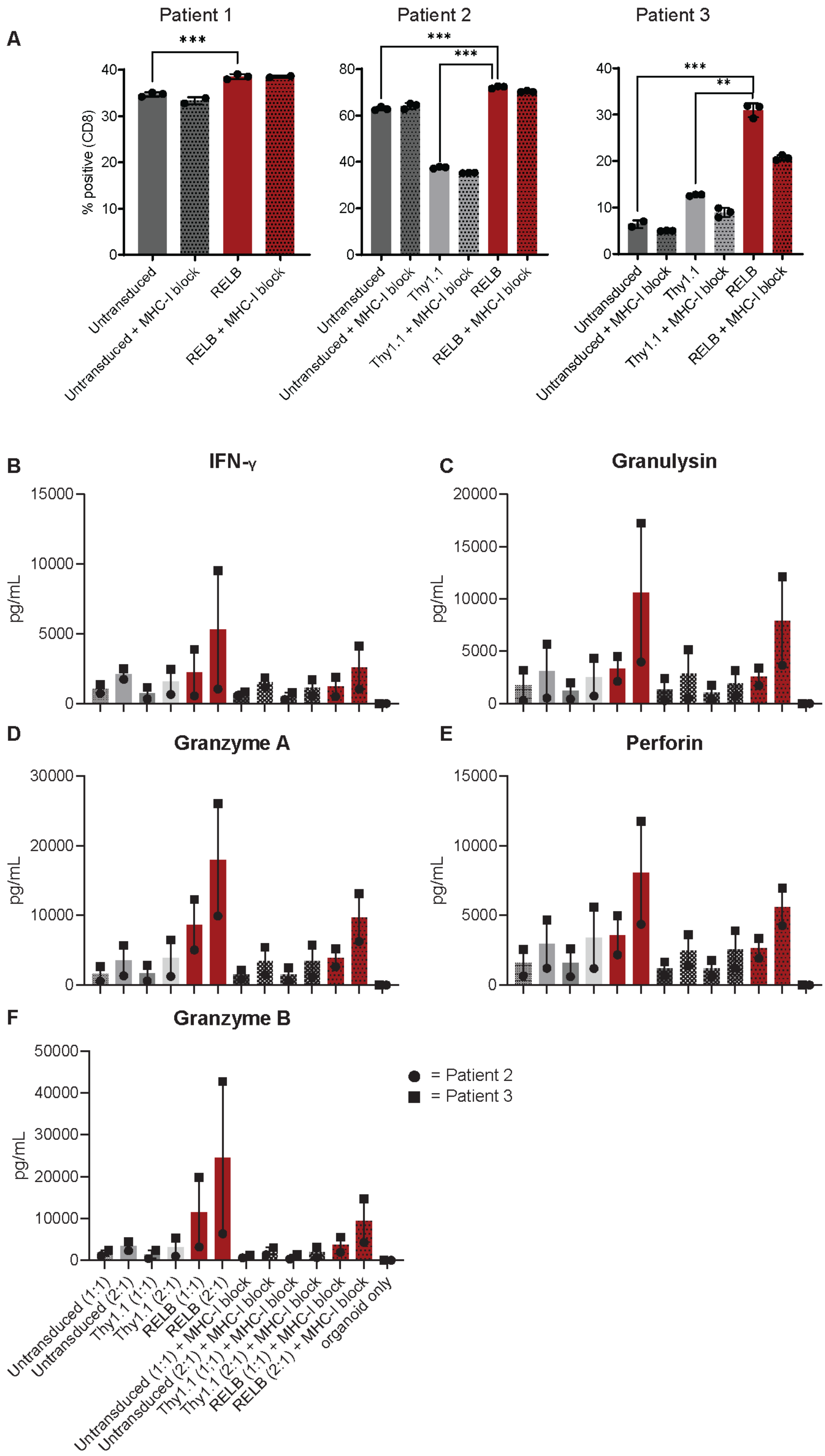

A
Patient 1
Patient 2
Patient 3
% positive (CD8)
***
**
Untransduced
Untransduced + MHC-I block
Thy1.1
Thy1.1 + MHC-I block
RELB
RELB + MHC-I block
B
IFN-γ
C
Granulysin
D
Granzyme A
E
Perforin
F
Granzyme B
pg/mL
= Patient 2
= Patient 3
Untransduced (1:1)
Untransduced (2:1)
Thy1.1 (1:1)
Thy1.1 (2:1)
RELB (1:1)
RELB (2:1)
Untransduced (1:1) + MHC-I block
Untransduced (2:1) + MHC-I block
Thy1.1 (1:1) + MHC-I block
Thy1.1 (2:1) + MHC-I block
RELB (1:1) + MHC-I block
RELB (2:1) + MHC-I block
organoid only

**Supplemental fig. 9: TIL+matched co-culture CD8+ staining and cytokine secretion flow cytometry analysis.** (A) Percent viable CD3+ CD8+ TIL events in co-culture samples for three patients. Error bars represent standard deviation. P. values were calculated using Welch's unpaired T test. * = < 0.05 , ** = < 0.01, and *** = <0.001 p. values. Patient 1 did not have control vector conditions due to insufficient growth. (B-E) Cytokine secretion flow cytometry quantification for patients 2 and 3 using bead-based Legendplex kit for (B) IFN-γ, (C) Granulysin, (D) Granzyme A, (E) Perforin, and (F) Granzyme B. Both 1:1 and 2:1 T cell:tumor cell co-culture ratios are shown.
